# Unilateral gene expression via maternally controlled zygotic recombination in *Drosophila*

**DOI:** 10.64898/2026.10.14.759140

**Authors:** Timothy A. Currier, David Tadres, Janina E. Troper, Yandan Wang, Thomas R. Clandinin

## Abstract

Structural and functional asymmetries are ubiquitous in bilaterians, but tools for unilateral genetic access have been lacking, a methodological gap that has limited mechanistic dissections of lateralized cellular functions. We developed the Maternally Controlled Zygotic Recombination (MCZR) system to overcome this limitation in *Drosophila*, co-opting the regulatory machinery of the maternal-to-zygotic transition to enable somatic recombination during early embryonic development. Targeting recombinase activity to this early window generates large somatic mosaics that frequently split across the anatomical midline. In conjunction with cell type-specific driver lines, this approach provides a reliable, scalable, and user-friendly platform for unilateral transgene expression. In proof-of-concept experiments, we apply our tool to the question of locomotor control, illustrating how MCZR can be used to unilaterally manipulate or monitor a cell type of interest. Overall, the MCZR system unlocks previously inaccessible experimental avenues across neuroscience, cell biology, and developmental biology.

## Introduction

Most animals, including mammals and arthropods, are members of the Bilaterian clade, possessing body plans characterized by mirror symmetry. This symmetry has a fundamental impact on many aspects of Bilaterian biology, including developmental tissue patterning and neural control of behavior^1,2^. However, despite the overall symmetry of the Bilaterian body plan, anatomical and functional asymmetries are present in many tissues^3^. How and why these breaks in symmetry occur are fundamental questions in evolutionary and developmental biology^4,5^. At the same time, functional asymmetries are widespread in the brain, with anatomically symmetric regions being involved in different sensory and motor processes across many Bilaterians^6,7,8,9,10,11,12,13,14,15^. Understanding the mechanisms of anatomical and functional asymmetry would benefit from tools that can unilaterally manipulate cellular function. However, individual cell types are almost always symmetrically distributed across the left and right sides of a tissue, meaning that even cell type-specific genetic reagents cannot ensure unilateral control. Addressing longstanding questions surrounding the genesis and functional consequences of asymmetry will therefore require new tools.

Prior efforts to overcome these limitations in *Drosophila* have relied on the generation of genetic mosaics to produce left-right expression asymmetries. In a classic approach, Hotta and Benzer utilized the spontaneous loss of a circularly permuted X chromosome to create mosaic “gynandromorph” flies during development^16,17^. Because this circular X chromosome is extremely unstable, it was typically lost within the first few embryonic divisions, resulting in large somatic mosaics that sometimes split across the midline. However, this same instability makes the experimental use of gynandromorphs challenging, and the resulting X monosomy reduces viability^16,18^. More recent strategies have employed targeted recombination to generate left-right expression asymmetries in specific cell types^19,20,21,22,23,24,25^. However, because these approaches were applied post-mitotically, they were incapable of creating large somatic mosaics, meaning that they scaled poorly as the population size of the targeted cell type increased beyond one or two cells per hemisphere^19^.

An alternative approach to generating large expression asymmetries would be to induce targeted recombination at very early stages of embryogenesis, when the left-right body plan is being established, and when each tissue is represented by a small number of progenitor cells. However, inducing somatic recombination at these early timepoints carries several technical challenges. For instance, the first 13 nuclear divisions in the developing fly are extremely rapid, and occur within a syncytial blastoderm^27^. During these divisions, homologous chromosomes do not pair^28^, potentially precluding mitotic recombination^29^. Furthermore, gene expression in the embryo is maternally controlled during these stages, with zygotic transcription only beginning more than two hours into development, during the mid-blastula transition (MBT). This maternal-to-zygotic transition (MZT) of gene regulation is a common developmental feature across the animal kingdom^30^ that complicates recombinase expression strategies. Together, the speed of these early divisions and the complexity of the genetic regulatory landscape suggest that recombination may not be feasible in the early fly embryo.

Here, we co-opt the genetics of maternal gene regulation and the MBT to create the Maternally Controlled Zygotic Recombination (MCZR) system, a set of genetic tools that target recombination to the earliest stages of fly development. We first characterize the timing of recombination events driven by a *smg-Flp* MCZR minigene, demonstrating that intra-chromosomal reactions, but not inter-chromosomal reactions, can occur during early nuclear divisions. We then use an analogous *smg-ФC31* MCZR construct to drive stochastic intra-chromosomal recombination during these early stages, producing large spatial mosaics that often split across the midline. In the brain, this strategy achieves reliable left-right expression asymmetries across a range of neuronal cell types. As a proof of concept, we apply our technique to the question of locomotor control, unilaterally manipulating and monitoring neural activity in a population of bilaterally projecting neurons. These data reveal functional asymmetries that are invisible to existing approaches, highlighting a role for these neurons in supporting interhemispheric coordination during locomotion. Overall, the MCZR system provides a robust, scalable, and user-friendly platform that opens new experimental avenues across neuroscience, cell biology, and developmental biology.

## Results

### Generating embryonic mosaics via Maternally Controlled Zygotic Recombination (MCZR)

As an initial step toward asymmetric, cell type-specific transgene expression, we first sought to induce recombination early in embryogenesis. At this stage, all tissues originate from a small number of nuclei, such that genomic recombination events will give rise to large somatic clones^17^. While integrases have been widely used for germ line transgenesis^31,32,33,34,35^, the ability of integrases and recombinases to catalyze intra- and inter-chromosomal rearrangements during early embryogenesis has been largely unexplored. We therefore sought to co-opt the regulatory machinery of the midblastula transition (MBT) to target recombinase activity to the earliest stages of embryogenesis.

To do this, we re-purposed the translational regulatory machinery of smaug (*smg*), an RNA-binding protein that is maternally deposited as mRNA into mature oocytes^36^. Smg mRNA is only translated upon egg activation, when an influx of calcium triggers protein synthesis^37^. We hypothesized that this regulatory system could be used to target recombinase activity to early embryonic development. To test this idea, we first engineered a MCZR minigene in which the open reading frame of Flp recombinase was flanked by the 5’ and 3’ regulatory sequences of smg, generating *smg-Flp* (Fig. 1A). With this approach, we reasoned that the translation of stockpiled Flp mRNA would be initiated shortly before fertilization, allowing recombinase activity to begin during the earliest zygotic stages^37,38^. Later in embryogenesis, during the MBT, proteins under smg regulatory control are actively degraded^37,38,39,40^. Thus, by ensuring that Flp was exclusively supplied through maternal deposition, we restricted recombinase activity to a very early developmental window (Fig. 1B).

**Figure 1.**
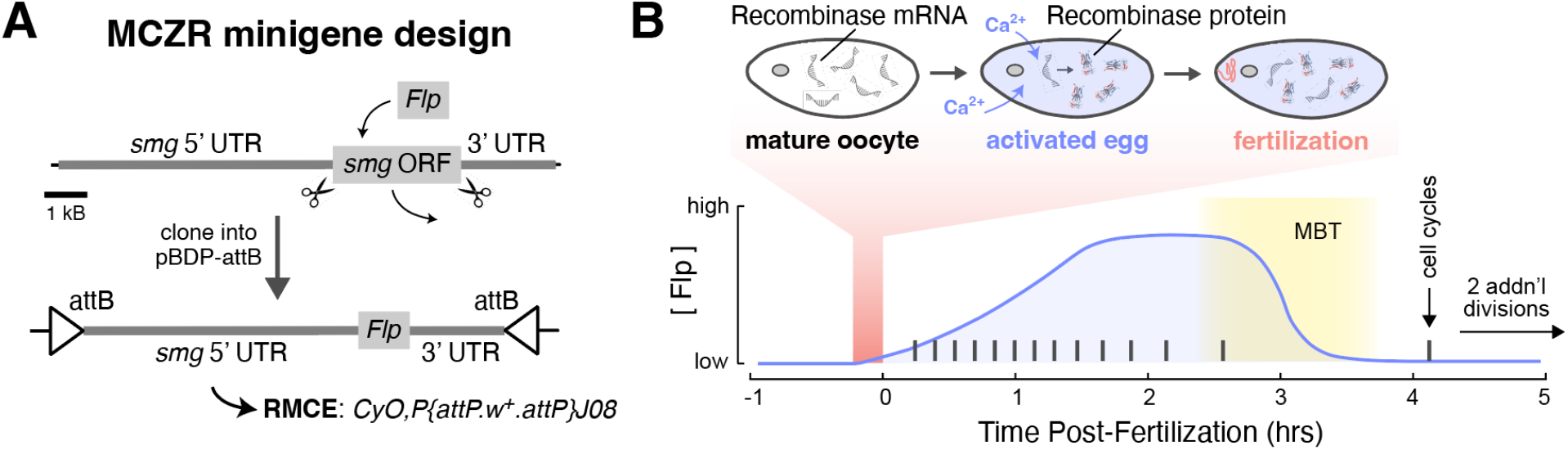
Generating embryonic mosaics via Maternally Controlled Zygotic Recombination (MCZR). **(A)** MCZR minigene design, with *smg-Flp* shown as an example. The *smg* locus was cloned^37^ and its open reading frame was digested out and replaced with *Flp* coding DNA. This entire 12 kB minigene was cloned into pBDP-attB and then inserted into a CyO,P{attP.w+.attP}J08 chromosome^35^ via recombination-mediated cassette exchange (RMCE). **(B)** Top: conceptual model of recombinase production in the oocyte under MCZR regulation. Smg UTRs prevent the translation of Flp mRNAs until calcium (Ca2+) floods the oocyte during egg activation^37^. Fertilization then immediately follows activation. Bottom: Conceptual model of MCZR recombinase concentration over developmental time, based on existing data^38,39^. Flp levels rise following egg activation and remain high until the midblastula transition (MBT), when maternal products are actively degraded. Each cell cycle is very short prior to the MBT. Methods used to assess this model of MCZR minigene function are detailed in Extended Data Fig. 1.

### Maternal smg-Flp drives recombination in the early embryo

Prior work on both Flp^41^ and serine integrases^42^ showed that intramolecular recombination events occur faster than intermolecular ones. Due to the rapidity of early nuclear divisions, we reasoned that a more efficient reaction would be more likely to occur in the early embryo (Extended Data Fig. 1A-D). We therefore turned to FlpOut^43,44^, a Flp-based reporter system with an intra-chromosomal recombination cassette, to ask whether recombination is possible in the early embryo. We assessed *smg-Flp*-driven recombination with a *UAS-FlpOut-GFP* cassette (MCZR-FlpOut), and imaged reporter expression in both adult and larval brains (Fig. 2A, B). We observed GFP expression in every MCZR-FlpOut fly that we imaged, confirming that recombination is indeed possible in the early embryo. Furthermore, GFP was expressed in nearly every neuron across the entire brain, consistent with early recombination (Extended Data Fig. 1B). Critically, GFP expression depended on the presence of *smg-Flp* in the maternal fly (Fig. 2A, B), confirming that the MCZR strategy can produce functional recombinase protein.

**Figure 2.**
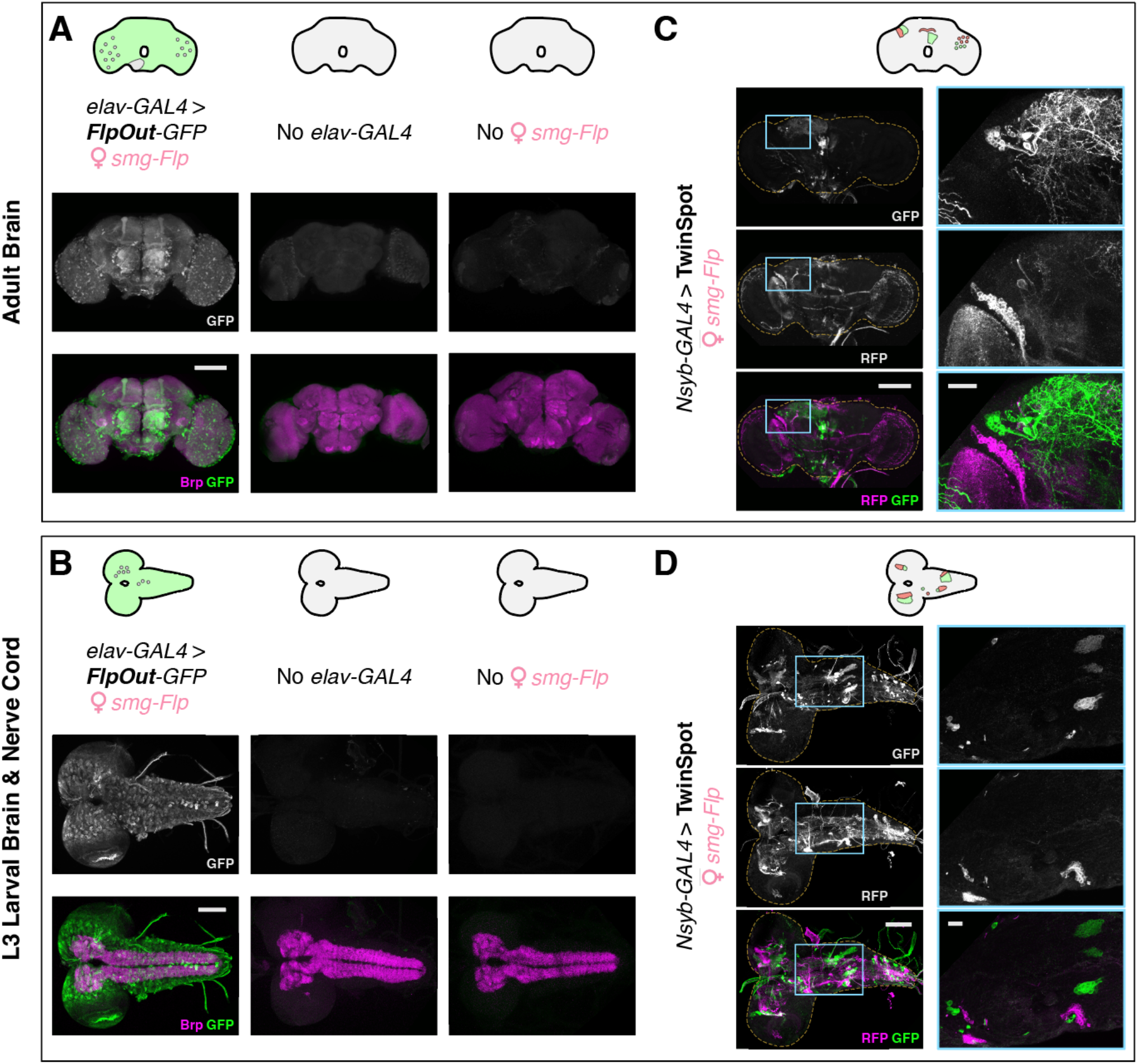
Maternal *smg-Flp* drives recombination in the early embryo. **(A**,**B)** Embryonic *smg-Flp* recombination driving FlpOut (i.e., MCZR-FlpOut) in adult brain **(A)** and L3 larval brain and nerve cord **(B)**. Top part of each panel contains schematic illustrations of expected results for each experiment, as in Extended Data Fig. 1. The bottom part of each panel shows confocal max intensity projections (MIPs) of immunostained tissue. Bruchpilot (Brp) counterstain shows neuropil extent. Scale bars: 100 μm. **(C**,**D)** Embryonic *smg-Flp* recombination driving TwinSpot MARCM (i.e., MCZR-TwinSpot) in adult brain **(C)** and L3 larval brain and nerve cord **(D)**. Top part of each panel: schematic illustrations of expected results. Bottom: confocal MIPs of immunostained tissue. For each set of images, dashed yellow lines indicate the tissue’s outer edge, and teal boxes in the left column (scale bar: 100 μm) indicate regions of magnification shown in the right column (scale bar: 20 μm). Magnified panels are partial MIPs to highlight specific clones. MCZR recombination in additional tissues are shown in Extended Data Fig. 2.

To better understand how *smg-Flp* functions in the early embryo, we next tested whether it can catalyze inter-chromosomal recombination reactions. Mosaic Analysis with a Repressible Cell Marker (MARCM^29^) reactions require recombination between homologous chromosomes, potentially constraining its efficiency^28,41^. We therefore expected that, if *smg-Flp* can drive any MARCM reactions at all during embryonic development, these events would lead to sporadic, small clones. To test this model, we used maternal *smg-Flp* to drive TwinSpot MARCM^45^ (MCZR-TwinSpot), which labels one daughter cell of a single division with RFP and the other with GFP (Extended Data Fig. 1C, D). These reporters persist, labeling both ensuing lineages and allowing us to visualize them in adult and larval brains (Fig. 2C, D). In contrast to the widespread GFP expression seen with MCZR-FlpOut, MCZR-TwinSpot produced small, sporadic GFP+ and RFP+ clones in almost all flies. This “patchy” expression pattern is consistent with the inter-chromosomal MARCM reaction being less efficient during the *smg-Flp* expression window (Extended Data Fig. 1D). Together, our FlpOut and TwinSpot results validate our model of *smg-Flp* function (Fig. 1B), and demonstrate that the MCZR strategy can catalyze both intra- and inter-chromosomal reactions in the embryo.

### Maternal smg-ФC31 can asymmetrize gene expression in a lineage-dependent manner

While MCZR-TwinSpot clones in the brain were small, they were almost always asymmetric across the midline, with each hemisphere displaying a unique clone distribution. This result suggested that embryonic recombination might indeed be a viable strategy for unilateral transgene expression. To reliably generate large-scale left-right asymmetries, we therefore sought a recombination cassette that (1) utilizes an intra-chromosomal reaction, like FlpOut, allowing efficient recombination at the earliest embryonic stages, and (2) produces two mutually exclusive and irreversible outcomes, like TwinSpot MARCM, to permanently encode the genetic asymmetry. Sparse Predictive Activity through Recombinase Competition (SPARC^46^) is a design that meets both of these criteria (Extended Data Fig. 1E). In SPARC, ΦC31 integrase selects stochastically between two attP target sites to irreversibly recombine with a single attB site, producing one of two mutually exclusive outcomes that either permit (“ON”) or prevent (“OFF”) GAL4-driven transgene expression.

By creating a *smg-ФC31* MCZR minigene, we sought to drive this stochastic intra-chromosomal recombination at the earliest stages of embryogenesis. After the SPARC reaction occurs in a given nucleus, its ON or OFF state will be inherited by its lineage (Extended Data Fig. 1F). Because early embryonic divisions define large spatial domains in the body plan, the timing of recombination should directly determine the size of somatic clones. In a pluripotent progenitor cell that gives rise to a tissue, a SPARC reaction would therefore produce uniform tissue expression (all ON or all OFF). As the left and right halves of the fly emerge from separate progenitors, independent SPARC reactions in those cells can produce different ON and OFF outcomes to generate left-right asymmetry. Progressively later recombination events would then produce progressively finer spatial patterns. In this way, MCZR-SPARC provides a strategy to convert early stochastic recombination events into spatially patterned asymmetry in any tissue, including the adult brain (Extended Data Fig. 3).

Using a pan-neuronal driver to visualize clones across the whole brain, we observed three outcomes of *smg-ФC31*-mediated SPARC recombination: fully ON (*N* = 22/122), fully OFF (*N* = 20/122), and left-right lateralized expression asymmetries (*N* = 80/122; Fig. 3A). Lateralized brains exhibited large asymmetric patches, which we defined as spanning at least a quarter of the central brain (*N* = 34/80), or small patches, spanning less than a quarter of the central brain (*N* = 46/80; Fig. 3B). Remarkably, we even observed complete hemisphere-scale asymmetry in a small number of flies (*N* = 3/122; Fig. 3A). These results suggest fly-to-fly variability in the precise timing of MCZR-SPARC recombination relative to the ongoing nuclear divisions in the embryo. Spatial variability was also observed, as asymmetric patch boundaries did not always align with the midline or antero-posterior and dorsal-ventral axes (Fig. 3A). These results demonstrate that early, maternally derived recombinase activity can efficiently generate large-scale, left-right expression asymmetry across many neuronal types.

**Figure 3.**
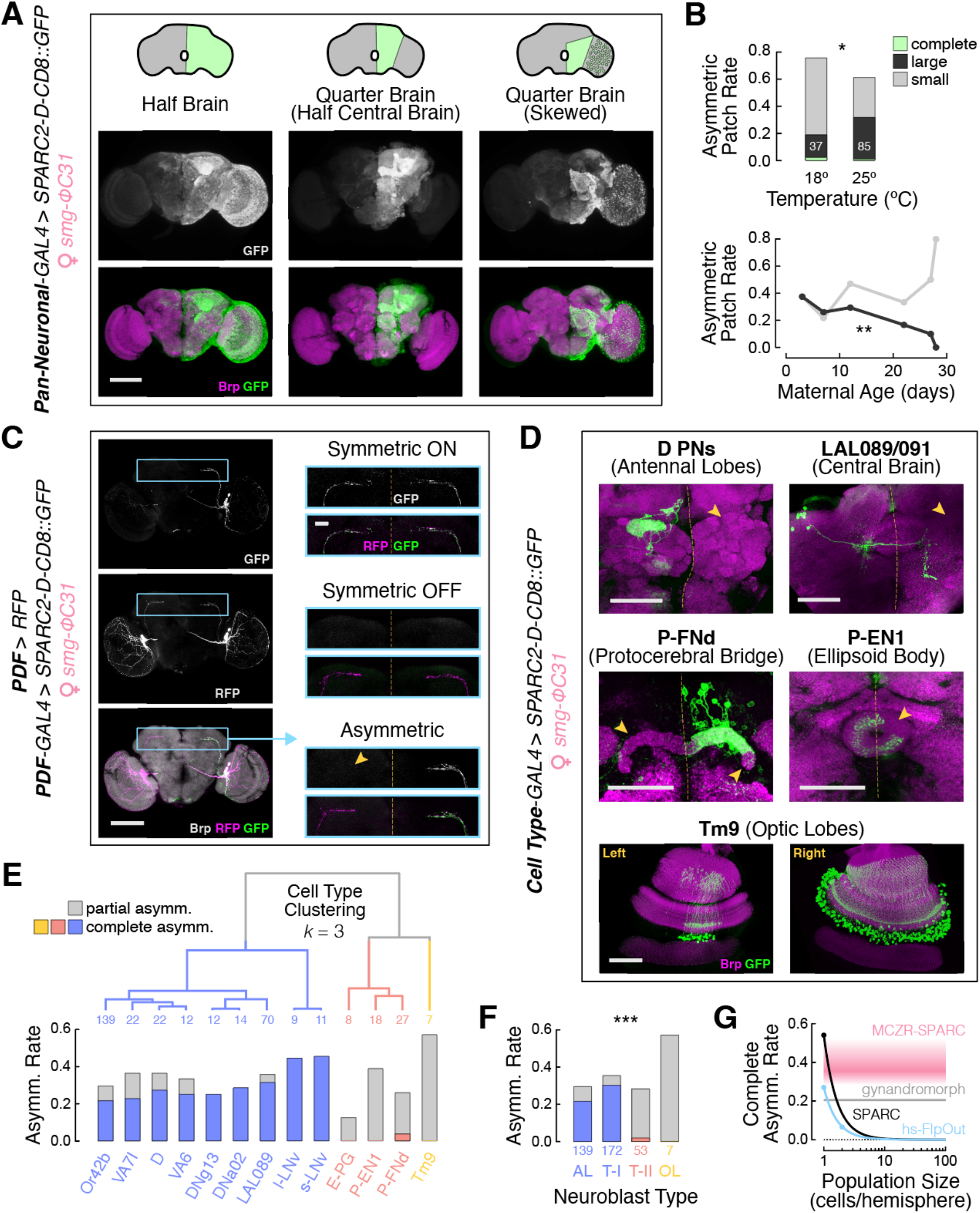
Maternal *smg-ФC31* can asymmetrize gene expression in a lineage-dependent manner. **(A)** Embryonic *smg-ФC31*-mediated SPARC recombination (i.e., MCZR-SPARC) in the adult brain of 3 example flies. Top: schematic representations of expression patterns, as in Extended Data Fig. 1. Bottom: confocal MIPs of immunostained adult brains. Brp counterstain shows neuropil extent. Scale bar: 100 μm. **(B)** Top: frequency of asymmetric GFP+ patches by patch size and temperature at the time of fertilization. Each fly is categorized by the size of its asymmetric GFP+ patch(es). Complete patches cover one entire brain hemisphere, large patches cover at least a quarter of the central brain, and small patches cover less than a quarter of the central brain. White numbers inside bars indicate the number flies in each group. *p* = 0.021 for large vs. small patch rate (Boschloo’s Exact). Bottom: frequency of asymmetric GFP+ patches by patch size and maternal age at the time of fertilization. Large patches: Spearman’s *ρ* = -0.94, *p* = 0.005; small patches: Spearman’s *ρ* = 0.71, *p* = 0.111. Group sizes for 3, 7, 12, 22, 27, and 28 days are 8, 23, 17, 12, 10, and 5, respectively. See also Extended Data Fig. 4 and Supplementary Table 1. **(C)** MCZR-SPARC recombination for an individual cell type (s-LNv). Left: an example full brain MIP. RFP is driven constitutively, while GFP depends on maternal *smg-ФC31* activity. Teal box indicates area of magnification. Scale bar: 100 μm. Right: magnified area for three different brains, one with symmetric GFP expression in s-LNv neurons, one with no expression, and one showing complete asymmetric expression. Asymmetric example brain is the same as shown on the left (teal arrow). Dashed yellow lines indicate the midlines in each image, and yellow arrowhead highlights missing GFP signal. Scale bar: 20 μm. **(D)** Examples of “complete” (top row) and “partial” (middle and bottom rows) asymmetry in different cell types. Each image is a MIP through the indicated brain regions. For Tm9 (bottom row), the left and right optic lobes of a single fly are shown as separate images. Scale bars: 50 μm. See also Supplementary Table 2. **(E)** *K*-means clustering of example cell types based on complete (colored bars) and partial (gray bars) asymmetry rates. At *k* = 3, cell types group approximately according to the neuroblast type from which they derive (yellow: Optic Lobe, red: Central Brain Type II, blue: Antennal and Central Brain Type I). *N* is shown at each leaf position in the dendrogram. **(F)** Complete and partial asymmetry rates by neuroblast type. While the overall rate of *any* asymmetry is consistent across neuroblast types (*χ*^2^ = 2.4, *p* = 0.48), *complete* asymmetry rates differ significantly (*χ*^2^ = 16.3, *p* < 0.001). Colors from (E). **(G)** Conceptual model of the scaling efficiency for different methods used to generate asymmetric mosaics in the fly brain. Pink band represents the observed floor (large patch rate, 28%) to theoretical ceiling (50%) of complete asymmetry rates for MCZR-SPARC across target population sizes. Darkest part of the band is the mean asymmetry rate across all tested cell types (33%). Lines represent population size scaling efficiency of post-mitotic ФC31 driving SPARC-D (black), post-mitotic hs-Flp driving FlpOut (blue), and the gynandromorph method^17^ (gray). Dots are real data points from other studies that used the post-mitotic SPARC^24^ and FlpOut methods^19^, and the corresponding lines are exponential extrapolations from those points.

To optimize the frequency of large-scale asymmetry, we next asked how different environmental and genetic factors influence the timing of recombination in the early embryo. We evaluated the rate and size of asymmetric clones across different temperatures, maternal ages, eclosion ranks, pan-neuronal driver lines, MCZR minigene insertion sites, and minigene dosages (Fig. 3B and Extended Data Fig. 4). Temperature and maternal age significantly influenced the rate of large asymmetric patches, with younger females and those mated at higher temperatures more frequently producing asymmetric offspring (Fig. 3B and Supplementary Table 1).

Given that the large asymmetric patches we observed appeared to fully encompass many cell types, we reasoned that complete left-right asymmetry would be even more common at the cell type level. As an example, we first examined asymmetry in sLNv neurons by driving *UAS-SPARC-D-CD8::GFP* in the presence of maternal *smg-ФC31* using *PDF-GAL4*. Each hemisphere contains four sLNv neurons^47^ – due to the stochastic nature of the SPARC reaction, many different expression patterns could emerge. On the one hand, symmetric outcomes could label all cells in both hemispheres, or no cells in neither hemisphere. Conversely, a completely asymmetric outcome would label four cells in one hemisphere, and none in the other hemisphere, while incompletely asymmetric outcomes would label variable numbers of cells on each side. We observed three of these outcomes: symmetric GFP ON, symmetric GFP OFF, and completely asymmetric GFP expression (Fig. 3C and Extended Data Fig. 3). Strikingly, 5/11 of flies displayed complete asymmetry, suggesting that MCZR-SPARC represents an efficient strategy for unilateral transgene expression at the cell type-level.

To systematically compare expression asymmetry across cell types, we examined *smg-ФC31* function across twelve driver lines spanning a diverse collection of neurons and glia (Fig. 3C-E, Supplementary Table 2, and Extended Data Fig. 2). Asymmetry was observed in all tested lines, at rates ranging from 12% to 57% depending on cell type, and 33% overall (*N* = 299), indicating that this strategy is broadly applicable. We performed *k*-means clustering based on each cell type’s rates of complete, partial, and no asymmetry (Fig. 3E). Remarkably, this unsupervised analysis segregated cell types approximately according to their developmental origins – cell types descended from central brain Type II neuroblasts formed one cluster, cell types descended from optic lobe neuroblasts formed another cluster, and a third larger cluster contained cell types descended from both antennal and central brain Type I neuroblasts. To explicitly ask whether developmental origin is predictive expression asymmetry, we grouped cell types according to their known neuroblast types and compared complete and partial asymmetry rates across groups (Fig. 3F). Consistent with our clustering analysis, optic lobe- and Type II-descended cell types showed significantly lower rates of complete asymmetry than Type I-descended cell types. Because optic lobe- or Type II-derived cell types are represented by single precursors much earlier in development than Type I-derived cell types^48,49,50^, complete asymetry for the former would require significantly earlier recombination events. Regardless, *smg-ФC31* maintains an experimentally useful rate of complete asymmetry for cell types with a wide range of population sizes, unlike prior methods (Fig. 3G). Overall, our results demonstrate that SPARC recombination driven by maternal *smg-ФC31* reliably produces scalable, lineage-dependent, and cell type-specific asymmetric transgene expression.

### Unilateral optogenetic activation uncovers otherwise obscured circuit functions

We next sought to demonstrate the versatility of the MCZR system through proof-of-concept experiments. Neural activity is highly lateralized during locomotor turns^15,51,52,53,54,55^, suggesting a degree of coordination between the two brain hemispheres. While connectome work^56,57^ has identified many bilaterally projecting cell types that could support such coordination, mechanistic studies have been difficult without an efficient system for unilateral monitoring and manipulation of candidate cell types^19,24,25^.

To address this gap, we used MCZR-SPARC to dissect the role of two Lateral Accessory Lobe (LAL) cell types – LAL089 and LAL091^56^, which are both labeled by the SS49952 split-GAL4^58^ driver line (Extended Data Fig. 5A). These bilaterally projecting cell types^59^, and other LAL neurons^20,25,60,61^, have recently been implicated in steering control. Previous work that bilaterally stimulated LAL089/091 neurons observed decreased groundspeed and a broadened angular velocity distribution^59^. However, unilateral perturbations of these neurons were not possible.

To better understand the function of LAL089/091 neurons, we placed individual flies in custom “FlyBowl” arenas^62^ and tracked their walking behavior using the Raspberry Pi Virtual Reality System (PiVR^63^). In these arenas, we prevented flies from walking upside-down by siliconizing the glass lid, allowing us to unambiguously identify left and right turns (Fig. 4A). Using maternal *smg-ФC31* and paternal *UAS-SPARC-D-CsChrimson* to express Chrimson in LAL089/091 neurons, we stimulated each fly with a series of 2 second pulses of red light while recording their walking trajectories (Fig. 4B). After behavioral testing, we dissected each experimental fly to determine whether Chrimson expression was unilateral, bilateral or absent (Fig. 4C, top). Unilateral expression patterns completely labeled either the left or the right hemi-population; the left hemi-population is the set of LAL089/091 neurons with their cell bodies in the left hemisphere, and vice versa for the right hemi-population (Extended Data Figure 5A).

**Figure 4.**
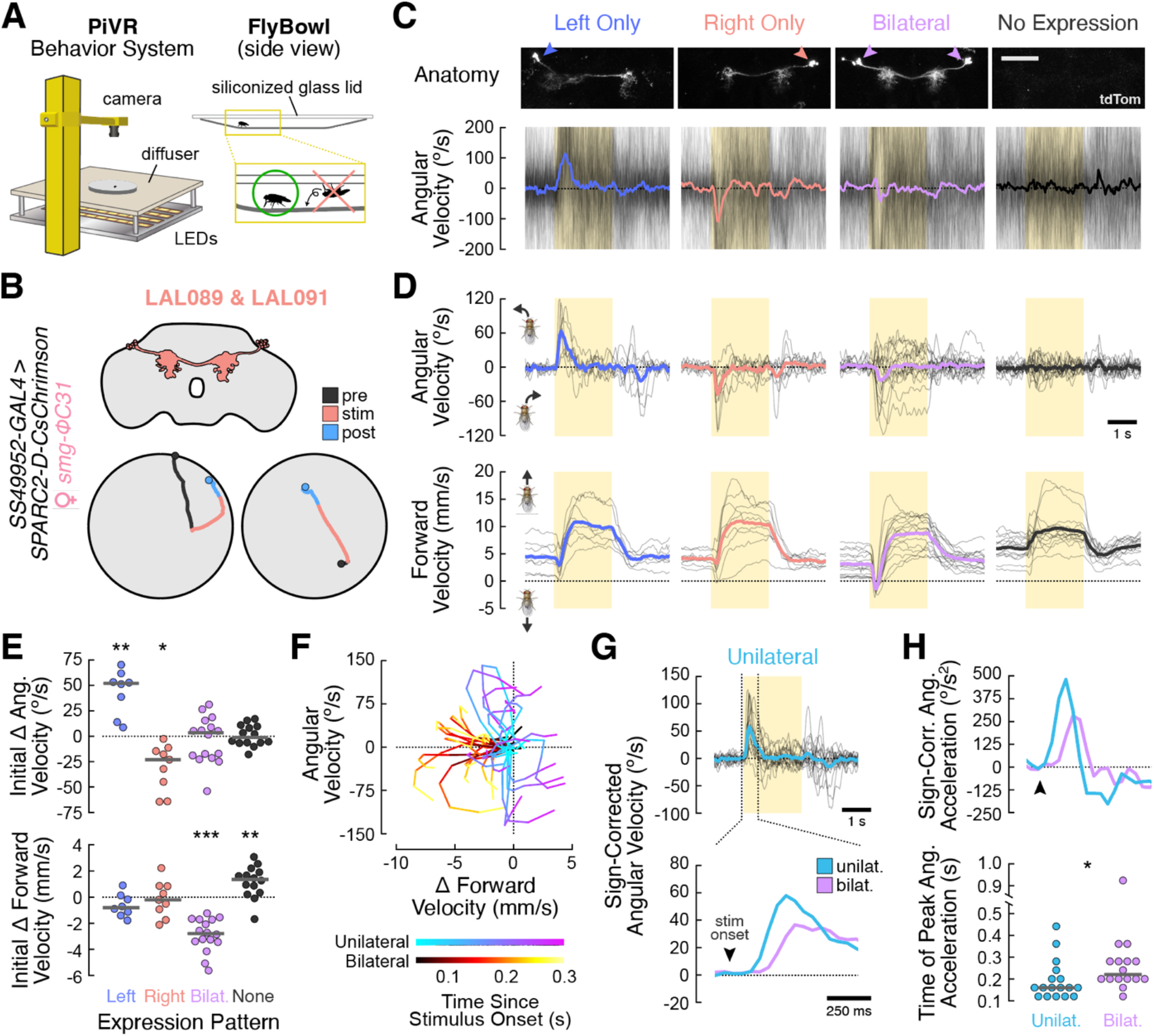
Unilateral optogenetic activation uncovers otherwise obscured circuit functions. **(A)** Monitoring unrestrained locomotor behavior using the PiVR system^63^. Single flies were placed in a bowl-shaped dish^62^, and their walking trajectories were tracked with a camera. The dish’s glass lid was coated in a siliconizing agent that rendered the surface too slippery for flies to walk on. **(B)** Proof-of-concept unilateral activation experiments. Top: cartoon depiction of the *SS49952-GAL4* expression pattern, which labels two connectomically defined cell types: LAL089 and LAL091^56,59^. See also Extended Data Fig. Bottom: single-trial example trajectories of two flies with MCZR-dependent CsChrimson expression in LAL089/091. Each of the plotted pre-stimulus (black), stimulus (red), and post-stimulus (blue) periods are 2 sec in duration. Black dots represent each fly’s starting position, blue dots represent each fly’s ending position, and the large gray circle indicates the approximate edge of the arena (see Methods). **(C)** Individual fly behavior. Top: *SS49952-GAL4* expression patterns driven by maternal *smg-ФC31* and *SPARC2-D-CsChrimson::tdTomato* for four individual flies. Each image shows a MIP through the lateral accessory lobe of brains imaged *ex vivo*, shortly after behavioral testing. Arrowheads point to the clusters of cell bodies used for categorizing expression patterns as left hemi-population only (blue), right hemi-population only (red), bilateral (purple), or no expression (black). Scale bar: 50 μm. Bottom: angular velocity over time for the same flies. Each thin black line is a single trail, with the yellow box indicating the 2 s optogenetic stimulation period. The cross-trial mean is plotted as a thick colored line for each fly. Positive values indicate leftward turns. **(D)** Group behavioral data. Cross-trial mean angular velocity (top) and mean forward velocity (bottom) is plotted for each fly as a thin black line, with flies grouped according to their expression patterns. Fly icons indicate directions of movement for positive vs. negative velocities. Cross-fly means are plotted as thick colored lines. Group sizes for left, right, bilateral, and no expression are 8, 9, 16, and 14, respectively. See also Extended Data Fig. 6. **(E)** Swarm plots of transient changes to angular velocity (top) and forward velocity (bottom) as a function of expression pattern. Each point is the cross-trial mean velocity change of a single fly from 0.05 to 0.3 s after optogenetic stimulus onset. Gray lines indicate cross-fly means for each group. Left hemisphere expression is associated with left turns (*p* = 0.003), right hemisphere expression with right turns (*p* = 0.011), bilateral expression with a decrease in forward walking (*p* < 0.001), and no expression with an increase in forward walking (*p* = 0.010). All effects by one-sample *t*-test with Bonferroni correction. **(F)** Phase diagram showing mean angular velocity vs. mean change in forward velocity over time for flies with unilateral (cool colors) or bilateral (warm colors) expression patterns. The unilateral group contains flies with either “left only” or “right only” expression patterns. Each line is a single fly, colored according to the time since optogenetic stimulus onset. See also Extended Data Fig. 6. **(G)** Top: Sign-corrected angular velocity for all unilateral flies, plotted as in (D). Each “left only” or “right only” fly’s mean angular velocity time course was multiplied by the sign of its initial turn after optogenetic stimulus onset. Dashed vertical lines indicate the period of magnification below. Bottom: cross-fly mean sign-corrected angular velocity for unilateral (teal) and bilateral (purple) flies. The time axis is expanded to highlight the different behavioral dynamics around optogenetic stimulus onset (black arrowhead). See also Extended Data Fig. 6. **(H)** Top: Mean sign-corrected angular acceleration for all unilateral (teal) and bilateral (purple) flies, plotted as in (G). Bottom: swarm plot showing the timing of each fly’s peak angular acceleration, relative to the onset of the optogenetic stimulus. The vertical axis is broken to more clearly show rank differences between groups (*p* = 0.017 by rank-sum test).

Using this approach, we identified 14 flies with no Chrimson labeling, 16 flies with bilateral expression, and 17 flies with unilateral expression. Flies expressing Chrimson only in the left hemi-population (*N* = 8) turned left at stimulus onset, while flies with expression only in the right hemi-population (*N* = 9) turned right (Fig. 4C-E). Flies with bilateral Chrimson expression chose a single turn direction on each trial, with some flies displaying cross-trial preferences toward one direction or the other. However, on average, bilateral flies showed neither a leftward nor a rightward turning preference, consistent with prior work^59^. Importantly, flies that did not express Chrimson in either hemi-population did not turn in response to stimulation. Together, these results reveal that LAL089/091 neuron activation drives ipsilateral turns.

We next examined forward velocity and found that all flies initiated forward runs in response to the optogenetic stimulus, including those with no Chrimson expression (Fig. 4D). These results suggest that red light stimulation itself increases forward velocity. To account for this innate red-light response, we corrected the forward velocity time course of each fly by subtracting the mean forward velocity time course of the “no expression” group. This analysis revealed a brief decrease in forward velocity at stimulus onset across the unilateral and bilateral expression groups (Extended Data Fig. 6A,B). While this decrease reflected slowed forward walking in unilateral flies, bilateral flies walked backwards during the first 300 ms after optogenetic stimulus onset (Fig. 4E, Extended Data Fig. 6C, and Supplementary Video 1).

Together, our analyses reveal different behavioral responses to unilateral vs. bilateral activation of LAL089/091: while unilateral flies slowed their forward walking and quickly began an ipsilateral turn, bilateral flies first walked backwards, and only then began turning (Fig. 4F). Furthermore, much of the turning induced by bilateral activation was executed during this backward walk. To quantify this difference in turn timing, we combined “left only” and “right only” groups by sign-correcting angular velocity (see Methods). We found that unilateral activation of LAL089/091 resulted in faster turns than bilateral activation (Fig. 4G,H and Extended Data Fig. 6D), consistent with a conflict between competing ipsilateral turning drives in bilateral flies.

Overall, our results demonstrate that LAL089/091 neurons are sufficient to drive ipsilateral turns, and suggest that these neurons could indeed be involved in interhemispheric coordination during locomotion. These experiments further highlight the utility of MCZR, which enabled a richer functional dissection than was previously possible with bilateral activation.

### Unilateral imaging of bilaterally projecting neurons provides deeper mechanistic insight

Many bilaterally projecting neurons are intertwined as they cross the midline and have overlapping projections in both hemispheres^56,64,65^. As a result, it is often impossible to spatially resolve the neural activity of a single hemi-population. Indeed, LAL089/091 neurons are a typical example of such cells. To measure the behaviorally correlated signals of each LAL089/091 hemi-population, we expressed a genetically encoded calcium indicator using *smg-ФC31* and *UAS-SPARC-D-GCaMP7f*. Flies walked on an air-supported spherical treadmill during 2-photon imaging, allowing for the simultaneous monitoring of locomotor activity and LAL089/091 calcium dynamics (Fig. 5A). Using manually defined ROIs, we first analyzed the left- and right-hemisphere neurites of LAL089/091 neurons in flies with bilateral GCaMP7f expression (Fig. 5B, Extended Data Fig. 5A). Left hemisphere neurites were positively correlated with right turns, while right hemisphere neurites were positively correlated with left turns (Fig. 5B,C). In contrast, neurites in both hemispheres were inhibited by forward walking and excited by backwards walking. When we computed linear filters to capture the relationship between locomotor behavior and neural activity, we found that the left and right hemisphere neurites had similar filters (Fig. 5C,D).

**Figure 5.**
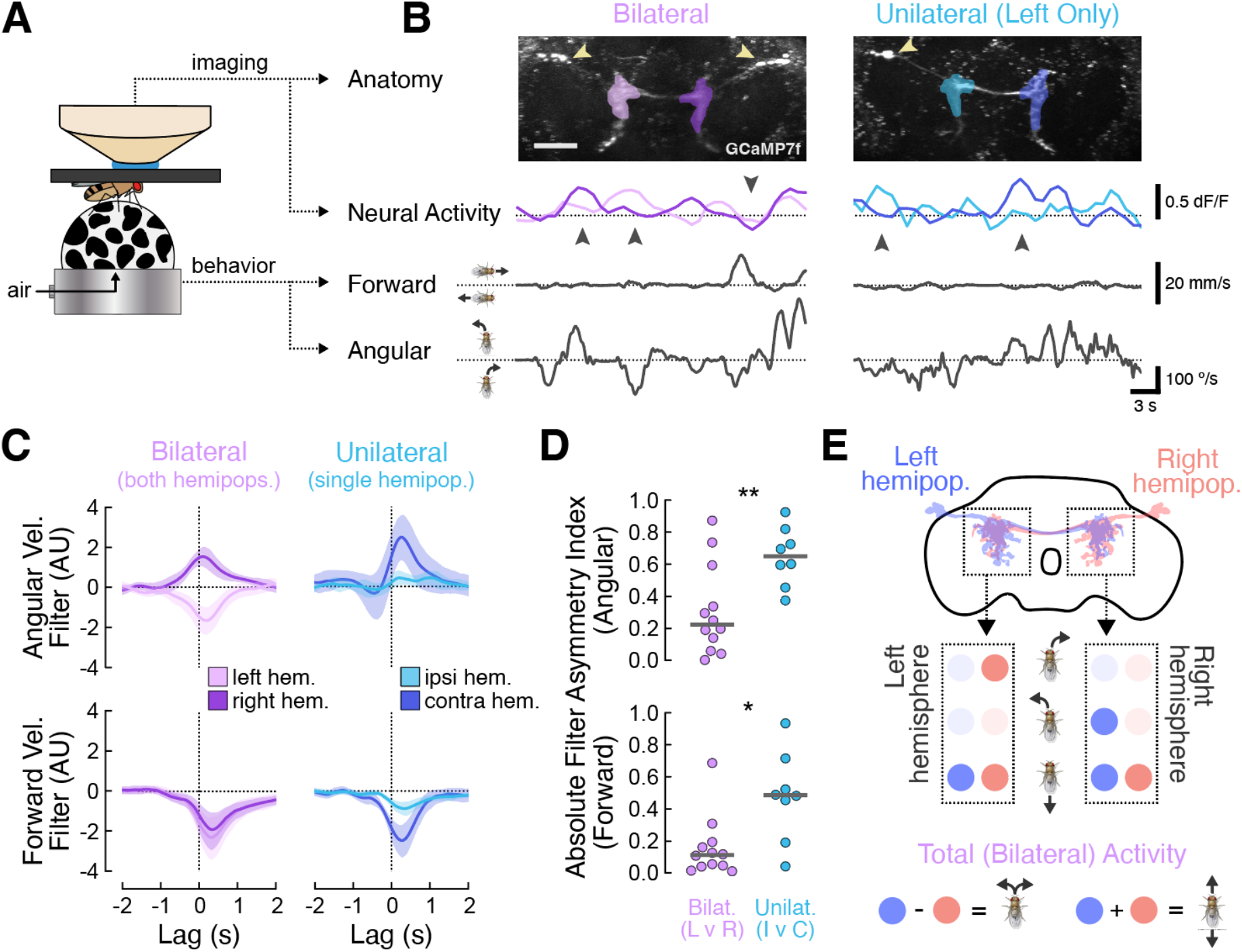
Unilateral imaging of bilaterally projecting neurons provides deeper mechanistic insight. **(A)** 2-photon imaging paradigm. Imaging and behavior data streams are collected simultaneously as each fly walks on an air-suspended ball (see Methods). **(B)** Top: example MIPs of two experimental flies, one with bilateral GCaMP7f expression in LAL089/091 neurons, and one with unilateral expression (left hemi-population only). The ROIs used for neural data collection are shown as colored overlays on each image. Yellow arrowheads indicate location of cell bodies. Scale bar: 50 μm. Bottom: 30 sec of simultaneous neural activity and movement recordings for the same flies. Neural activity traces are colored according to their respective ROIs. Upward arrows highlight angular velocity-correlated activity, while the downward arrow highlights forward velocity-correlated activity. **(C)** Linear filters that describe the relationship between LAL089/091 activity and either angular velocity (top) or forward velocity (bottom). Solid lines are the mean filter, with shaded regions illustrating the 90% CI. Left: For bilateral flies, filters are independently calculated for neurites in the left and right hemispheres. Because LAL089/091 neurons have bilaterally overlapping projections, these filters represent the summed activity of both hemi-populations. Positive angular velocity filter values represent excitatory responses to leftward turns. Right: for unilateral flies, filters are independently calculated for neurites either ipsilateral or contralateral to the cell bodies. These filters necessarily represent the activity of a single hemi-population. Here, positive angular velocity filter values represent excitatory responses to ipsilateral turns (relative to the side of the cell bodies). **(D)** Swarm plots of filter asymmetry index for angular (top) and forward velocity filters (bottom). Each point represents a single fly. Bilateral filters are more symmetric across the two hemispheres (lower asymmetry values), meaning that unilateral imaging provides a significant amount of additional information beyond what can be learned with bilateral imaging (angular velocity: *U* = 81, *p* = 0.009; forward velocity: *U* = 80, *p* = 0.012). **(E)** Summary of behavior and imaging results. Top: the activity of the left and right hemisphere neurites of each LAL089/091 hemi-population is represented as a pair of colored dots, with deeper saturation indicating higher activity levels. During rightward turns, only the left hemisphere neurites of the right hemi-population are active. During leftward turns, only the right hemisphere neurites of the left hemi-population are active. All neurites from both hemi-populations are active during backward walking. See also Extended Data Fig. 7. Bottom: the total difference in activity between the left- and right-hemi-populations is a continuous, signed readout of angular velocity. The total summed activity across the left- and right-hemi-populations is a continuous, signed readout of translational velocity.

Because these filters reflect mixed signals from the left and right hemi-populations, these results cannot distinguish between two distinct models by which these neurons encode motor information. In the first model, ipsilateral turns could be encoded by neurites contralateral to the cell body – for example, left hemi-population neurons would represent left turns with activity in the right hemisphere neurites (Fig. 5E, Extended Data Fig. 7). Alternatively, in a second model, contralateral turns could be encoded by LAL089/091 neurites ipsilateral to the cell body – here, left hemi-population neurons represent right turns with activity in the left hemisphere neurites (Extended Data Fig. 7). To differentiate between these models, we calculated analogous filters for flies with unilateral GCaMP7f expression. In contrast to bilateral flies, we found substantial differences between the locomotor signals encoded in the two hemispheres (Fig. 5B-D). Specifically, LAL089/091 neurites ipsilateral to their cell bodies carried essentially no information about angular velocity, and very little information about forward velocity. In contrast, contralateral neurites were both robustly activated by contralateral turns and strongly inhibited by forward walking (Fig. 5C). These results imply that the symmetric activity patterns seen for each hemisphere in bilateral flies are almost exclusively originating from contralateral LAL089/091 neurons, supporting the first model (Fig. 5E).

Our data show that each LAL089/091 hemi-population specifically sends ipsilateral turning signals to the contralateral hemisphere (Fig. 5E). In further support of this model, a connectomic analysis revealed that nearly all of the presynaptic sites (output synapses) in these cells are located contralateral to the cell body (Extended Data Fig. 5B). This organization suggests a directionality to the information flow in LAL089/091, consistent with the functional compartmentalization observed in our calcium imaging experiments. Furthermore, the primary input and output cell types to LAL089/091 are visual projection neurons in the LAL and Anterior Optic Tubercle (Supplementary Table 3). Because this latter region encodes navigation-relevant visual cues^66^, these synaptic partners suggest that LAL089/091 neurons may integrate interhemispheric visuomotor signals to guide navigation. Overall, our results demonstrate the deeper mechanistic intuition that can be achieved with MCZR, offering clear evidence that LAL089/091 supports interhemispheric coordination during locomotion.

### Additional experimental use cases for the MCZR system

To highlight the experimental versatility of the MCZR system, we performed two additional proof-of-concept studies. To this point, our unilateral manipulations had been limited to existing SPARC constructs. To extend the system to any arbitrary effector, we combined *smg-ФC31* with SPARC3-OUT^67^, thereby enabling unilateral expression of GAL80, an inhibitor of GAL4. To test this combination, we used the *PDF-GAL4* line to drive a *UAS-6XGFP* reporter in the presence of smg-ФC31 and SPARC3-OUT (Fig. 6A). We found that SPARC3-OUT produced asymmetric GFP expression, demonstrating that the MCZR system can asymmetrize any UAS-based construct.

**Figure 6.**
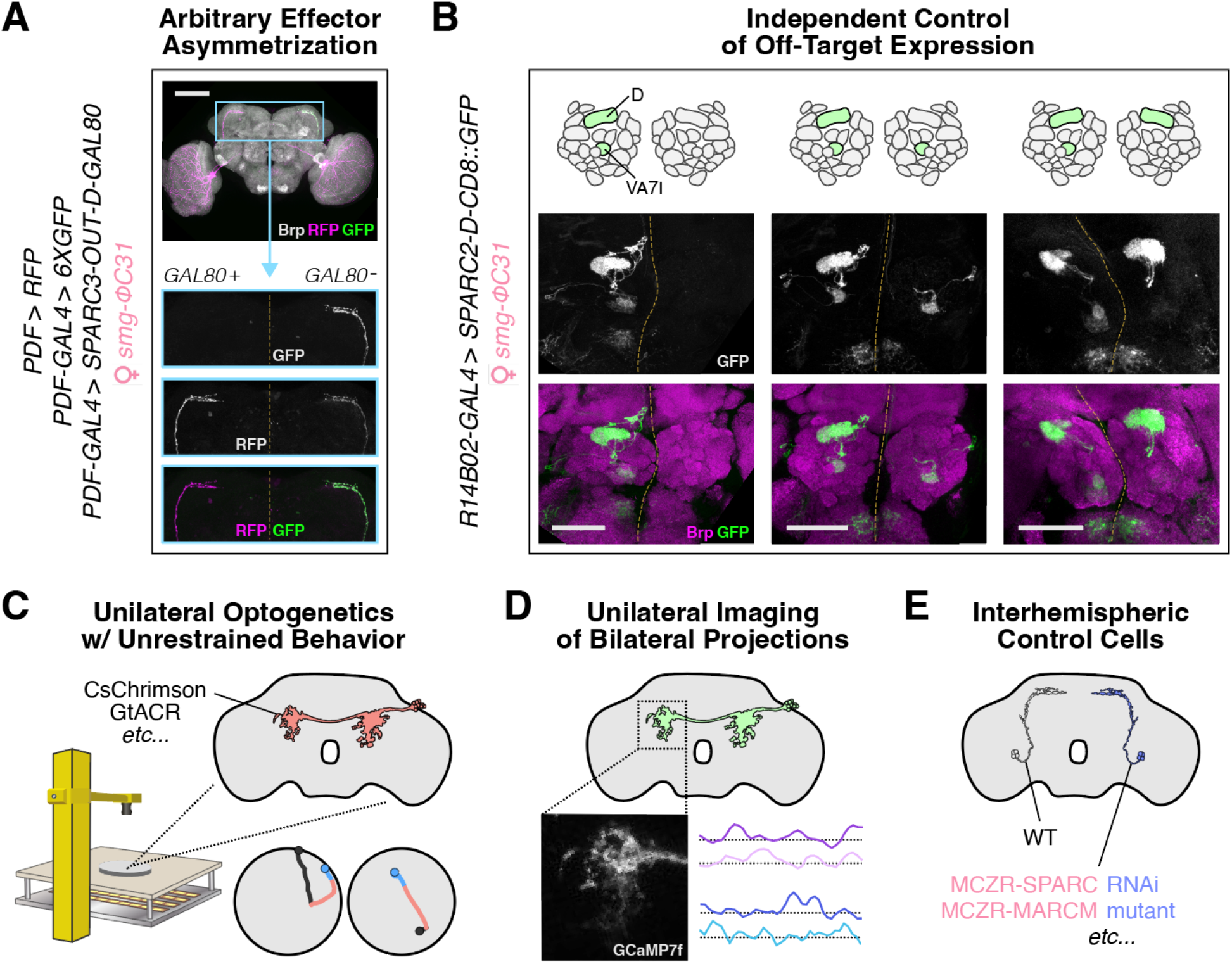
Experimental use cases for the MCZR system. **(A)** Asymmetric expression of any arbitrary effector via *SPARC3-D-OUT-GAL80*^67^. Instead of directly driving GFP, MCZR-SPARC recombination produces GAL80, which inhibits GAL4 and prevents constitutive GFP expression. GAL4-independent RFP expression is shown as a control. Top: example MIP of SPARC3 recombination in s-LNv neurons, as in Fig. 3C. Teal box indicates region of magnification below. Bottom: magnified images of the same brain. Dashed yellow lines indicate the midline. Scale bar: 100 μm. **(B)** Independent control of off-target expression. Top: schematic illustration of GFP expression from three example R14B02-GAL4 flies, which normally drives bilateral expression in two olfactory PN populations: D and VA7I. Bottom: corresponding antennal lobe MIPs. Scale bars: 50 μm. **(C)** Unilateral optogenetics during unrestrained behavior, as in Fig. 4. **(D)** Unilateral imaging of neurons with bilaterally overlapping projections, as in Fig. 5. **(E)** Within-fly interhemispheric control cells for any arbitrary Flp- or ФC31-dependent manipulation, as in (A).

Finally, we explored how MCZR-SPARC would behave when combined with driver lines that label more than one cell type. Due to the stochastic nature of mosaics generated with MCZR-SPARC, we reasoned that effector gene expression in each hemisphere should be independent for each cell type. In the context of behavioral studies, such independent expression could be used to control for off-target activation or silencing phenotypes. We utilized *R14B02-GAL4*, which labels two populations of olfactory projection neurons (PNs) – D and VA7I – and drove GFP expression downstream of *smg-ФC31* with *UAS-SPARC-D-CD8::GFP* (Fig. 6B). Across flies, we observed GFP expression in all possible combinations of PN types and hemispheres. These results demonstrate that MCZR-SPARC can separate individual cell types in complex driver lines. Collectively, our proof-of-concept studies demonstrate the versatility, scalability, and user-friendliness of the MCZR system.

## Discussion

### Unlocking recombination in the early embryo

We developed the MCZR system with the goal of inducing somatic recombination in the early embryo, a developmental window in which recombination has been thought to be challenging. Indeed, while recombination reactions have been successful in both the oocyte^68^ and post-MBT in the embryo^29,69^, the roughly 3-hour period between fertilization and cellularization has remained inaccessible. We overcame this limitation by co-opting the fly’s native MZT machinery. Many genes, including *smg*, are maternally deposited into mature oocytes as mRNA, translated upon egg activation, and actively degraded during the MBT^37,38,39,40^. We leveraged this regulatory time course – a perfect match to the embryonic recombination temporal targeting gap (Fig. 1) – to generate functional recombinase protein in the very early embryo.

While the precise timing of zygotic recombination with MCZR – inferred through clone sizes – varied considerably from fly to fly, we found that the overall distribution of reaction times was sensitive to the geometry of each recombination reaction. Specifically, intra-chromosomal reactions like MCZR-FlpOut and MCZR-SPARC occurred earlier in development than inter-chromosomal reactions like MCZR-TwinSpot (Fig. 2). These results are consistent with inter-chromosomal mechanisms being intrinsically less efficient than their intra-chromosomal counterparts^41,42^, and that homologous chromosomes do not pair in the early embryo^28^. These factors together likely account for the paucity of somatic clone studies in the early embryo.

With MCZR-SPARC, we found that higher temperature and lower maternal age produced flies with larger patches of asymmetric expression, consistent with earlier recombination events (Fig. 3B and Extended Data Figs. 1 and 4). Temperature and maternal age influence the rate of egg laying^70^, suggesting that the speed of maternal reproductive function might be positively correlated with recombination efficiency in the early embryo. This result might appear counter-intuitive – if eggs move more quickly through the reproductive tract, the interval between activation and fertilization is expected to be shorter, providing less time for recombinase translation before embryonic development begins (Fig. 1). However, warmer temperatures and younger maternal flies may also optimize metabolic conditions, potentially allowing recombinase translation or enzymatic function to outpace increased reproductive motility. Ultimately, this “race” between sufficient recombinase production and fertilization appears well balanced across a range of conditions, as most of the environmental and genetic factors we tested did not modify recombination efficiency.

However, the specific developmental window over which *smg-ФC31* recombination occurs ultimately limits its utility for generating clones at the largest spatial scales, up to and including the whole-animal level. Future studies might accelerate reaction timing by employing alternative regulatory frameworks. Compared to *smg, Cup* and *ME31B* are translated in earlier, shorter pulses at the beginning of embryogenesis^38^, suggesting that Cup or ME31B MCZR minigenes might outperform *smg-ФC31* for creating large asymmetric clones. Similarly, work in other recombinase systems has shown that the distance between recognition sites can dramatically influence recombination rate^42^, suggesting that modifications to SPARC cassettes could tune reaction timing. Because the MZT is a highly conserved developmental feature across the animal kingdom^30^, it is also likely that the MCZR strategy can be translated to other model systems. Regardless, the availability of both Flp- and ΦC31-mediated somatic mosaics opens new possibilities for exploring non-autonomous interactions amongst nuclei in early development, and paves the way for a next generation of embryological studies.

### A robust, scalable, and user-friendly approach to unilateral transgene expression

As a specific use-case for the MCZR system, we performed *smg-ФC31*-mediated embryonic SPARC reactions to generate large-scale somatic clones that frequently split across the anatomical midline (Fig. 3A). Across cell types and brain regions, about one-third of flies displayed complete unilateral expression patterns, a vast improvement over existing methods (Fig. 3G). For example, post-mitotic methods utilizing ФC31^21,23,24,25^ or heat-shock Flp^19,26^ achieve high rates of complete asymmetry only for cell types with a single neuron per brain hemisphere. Complete asymmetry using these prior methods is exponentially less likely as population size increases, becoming experimentally impractical with more than two cells per hemisphere. Conversely, the experimentally challenging gynandromorph approach scales well across population sizes, but produces asymmetric flies at a lower overall rate than the observed floor achieved with *smg-ФC31*^17^. Thus, MCZR-SPARC uniquely combines high efficiency with scalability, providing a robust new method for driving asymmetric gene expression across a wide range of cell types.

Our experiments in LAL089/091 provide a clear demonstration that MCZR will be an essential new tool for circuit-level dissections of locomotor control. We show that MCZR is a user-friendly system for performing unilateral stimulation and imaging experiments, that intrinsically incorporates all relevant controls. Taking unilateral optogenetic activation as an example, bilateral and “no expression” experiments are necessary statistical comparators^71^. With MCZR-SPARC, the same experimental cross that generates unilateral flies will also produce these exact controls, in approximately equal proportions (Fig. 4 and Extended Data Fig. 3). Furthermore, bilateral activation and imaging of LAL089/091 produced results that obscured the function of these cells, highlighting the vital importance of unilateral manipulations. In the absence of unilateral stimulation experiments, we might have concluded that LAL089/091 possess a functional role similar to that of Moonwalker Descending Neurons (MDNs^19,26^), which induce backward walking during bilateral activation. However, unilateral activation of LAL089/091 produced forward turns, while unilateral activation of MDNs produced backward turns^19^ (Fig. 4 and Extended Data Fig. 6). These results highlight how experiments with MCZR-SPARC can provide deeper functional descriptions and allow researchers to avoid potentially misleading conclusions.

Beyond imaging and activation, combining *smg-ФC31* with SPARC3-OUT allows for the unilateral expression of any arbitrary construct. For example, unilateral knockdown or overexpression of genes of interest can reveal cell biological or circuit-level consequences, with the added benefit of unperturbed “control” cells existing in the opposite hemisphere of the same experimental animal. Such experiments would eliminate many concerns surrounding common confounding variables, including genetic background, individual animal experience, or circadian phase.

Finally, we show that MCZR-SPARC can distinguish between on- and off-target effects when using driver lines that label more than one cell type. Here the “incompleteness” of asymmetric gene expression is an advantage, allowing an experimenter to localize specific stimulation effects to discrete cell types within an otherwise complex driver line. This ability of MCZR to control for off-target effects dramatically expands the space of driver lines that can be used for behavioral screens by circumventing the need for clean, cell-type specific lines, which may not be available for a cell type of interest.

Together, our experiments demonstrate the robustness, versatility, and user-friendliness of the MCZR system. By enabling recombination in the early embryo, and by allowing for efficient, cell type-specific unilateral transgene expression, MCZR can open new experimental avenues for cell biology, developmental biology, and neuroscience.

## Acknowledgements

The authors would like to thank Howard Lipshitz for generously providing CaSpeR4-SGS plasmid, Liqun Luo and Katherine Nagel for sharing fly stocks, and Jason Millington for help with immunohistochemistry materials. Thanks as well to Claude Desplan, Yvette Fisher, and members of the Clandinin Lab for valuable discussions and feedback on the manuscript. This work was supported by a Kirschstein National Research Service Award (F32EY035135, TAC), by a Swiss National Science Foundation Postdoc.Mobility Fellowship (222161, DT), by a Sapp Family Bio-X Undergraduate Summer Research Fellowship (JET), by the Stanford Vision Core (P30EY026877, TRC), and by Investigator Funds from BioHub San Francisco (TRC).

## Author Contributions

Conceptualization, TAC and TRC; Methodology, TAC, DT, and TRC; Investigation, TAC, DT, JET, and YW; Software, TAC, DT, and JET; Formal Analysis, TAC, DT, and JET; Visualization, TAC and DT; Writing – Original Draft, TAC, DT, and JET; Writing – Review & Editing, TAC, DT, JET, YW, and TRC; Funding Acquisition, TAC, DT, JET, and TRC; Supervision, TRC. TAC, DT, and JET contributed equally to this work and therefore possess equal rights to list themselves first in bibliographic documents.

## Methods

### Fly stocks and maintenance

Unless otherwise noted, all fly stocks were maintained at 18º C and 60% relative humidity, on a strict 12-hour light/dark cycle, on standard molasses food. All crosses were set up at 25º C. Supplementary Table 4 lists the exact genotypes of flies shown in each Figure, as well as the source of each stock. All newly generated fly strains will be made publicly available via the Bloomington Drosophila Stock Center.

While testing different environmental and genetic factors for effects on the rate of MCZR-SPARC expression asymmetry (Fig. 3 and Extended Data Fig. 4), virgin females were collected shortly after eclosion at 25° C and subsequently maintained at 18° C until subsequent mating at 25° C. Following mating, experimental progeny were reared at 25° C and collected over the course of 10 days. The only exception to this protocol was made when testing for the effect of temperature, when virgin females were mated and housed at 18º C for four days. Because the relevant recombination reactions occur early in development, rearing experimental progeny at 18º for longer periods of time was deemed unnecessary. As a result, progeny were transferred after these four days to 25º C, then collected over the course of 10 days.

While testing the influence of eclosion rank, single females were crossed to single males and experimental progeny were collected twice per day for 2 days. Additional collections were performed over the ensuing week at a reduced frequency to obtain later-born age groups. Because no continuous relationship between eclosion rank and asymmetry rate was observed, flies were grouped into “first day” and “after first day” eclosion timing groups for statistical comparison (Extended Data Fig. 4D).

For behavioral experiments, flies were raised at 25º C on standard molasses food containing 0.1 mM all-*trans*-retinal (ATR) (Sigma-Aldrich, R2500-500MG). All vials containing ATR food were protected from light by wrapping them in aluminum foil. Crosses were initially set up on ATR food, and experimental progeny were flipped onto fresh ATR food 3 days prior to behavioral testing. After this flip, flies were moved to new incubators with staggered light-dark cycles, and left for 3 days. This protocol ensured that, on the day of behavioral testing, all experiments were conducted between ZT23 and ZT02 (1 hour before subjective dawn through 2 hours after).

### MCZR molecular genetics

All sequence analysis and construct design was performed in Geneious (Dotmatics). CaSpeR4-SGS plasmid^37^ (*smg5’UTR-GFP-smg3’UTR*; kind gift of Howard Lipshitz) containing a GFP open reading frame flanked by *smg* UTRs was verified for sequence integrity via NotI-HF and NheI-HF restriction analysis (New England Biolabs, #R3189S and #R3131S). CaSpeR4-SGS was transformed into chemically competent *E. coli* (OneShot Top10, Invitrogen). Due to the large size of the SGS insert (>9000 bp), many transformed colonies lost SGS while retaining Ampicillin resistance. Smaller, slower growing colonies were therefore selected from among transformants, and were again verified via restriction analysis.

The 9502 bp product of NotI-HF digestion, which contained the SGS construct, was cloned into pBDP, producing pBDP-SGS. From this step onward, all subsequent plasmid products were verified via sequencing (Sequetech). The following primers were used to detect the presence of the *smg* 3’ UTR: smg3’_F1 (AACCGTAACTGCACCAGAGG), smg3’_R1 (AGGAGGAATGCGTTCACTGG), smg3’_F2 (ACATCGCTTCAACAACGTGC), and smg3’_R2 (CGCAGATTCTTACGCTTGGC). For the *smg* 5’ UTR, these primers were used: smg5’_F1 (GAATATGCCAAGCGCCTGTG), smg5’_R1 (CGCCGTTGCTGTTGTTGTTA), smg5’_F2 (GCAGCATCGCATTCCTCTTG), and smg5’_R2 (CAGCTCGTTTTGATACGCGG). Primers for detecting recombinase cDNA and PCR materials are detailed below (“Tissue collection and genotyping”).

Using GenEZ cloning (Genscript), the GFP open reading frame in pBDP-SGS was replaced with ΦC31 cDNA, sourced from pJFRC-Nsyb-IVS-ФC31 (Addgene #133868). The resulting pBDP-SΦS was injected into various P[acman]^32^ embryos for ΦC31-mediated genomic integration (BestGene). These injections produced the first generation of *smg-ФC31* MCZR lines: *yw; PBac{y[+]w[+mc]smg-ФC31}VK11, yw; PBac{y[+]w[+mc]smg-ФC31}VK18*, and *yw; PBac{y[+]w[+mc]smg-ФC31}VK33*.

These strains were marked with mini-white and could not be selected against during experimental crosses, prompting us to create a second generation of *smg-ФC31* MCZR lines that had the core SΦS minigene inserted into balancer chromosomes. We removed mini-white from the pBDP backbone via AscI digestion (New England Biolabs, #R0558S) and added a second, inverted attB site 56 bp 5’ of SΦS via GenEZ cloning, yielding pBDP(del: w[+mc])-SΦS-attB, which was injected into Bateman strains^35^ for recombination-mediated cassette exchange (RMCE) onto balancer chromosomes. The 56 bp between the inverted attB site and the *smg* 5’ UTR acted as a linker and was therefore integrated during RMCE, and had the following sequence: GTTTAAACGGCCGGCCAGATCTGGTACCGGATCCCCGCGGCCGCTGATCCTCTAGA. These injections produced the second generation of *smg-ФC31* MCZR lines: y*w; CyO,P{smg-ФC31}J08* and *w; TM3,Sb,Ser,P{smg-ФC31}FS10*. The data in Fig. 3 and Extended Data Fig. 4 were collected with a combination of first and second generation *smg-ФC31* lines, while all other experiments exclusively used the second generation.

Finally, an analogous pBDP(del: w[+mc])-SFS-attB construct was created via GenEZ cloning by replacing the ΦC31 cDNA in pBDP(del: w[+mc])-SΦS-attB with Flp cDNA, sourced from pHD-UAS-SPARC-D-Flp^46^. pBDP(del: w[+mc])-SFS-attB was injected into the same Bateman lines to create the following *smg-Flp* MCZR lines via RMCE: y*w; CyO,P{smg-Flp}J08* and *w; TM3,Sb,Ser,P{smg-Flp}FS10*.

### Tissue collection and genotyping

Unless otherwise specified, flies were aged 3-7 days post-eclosion (DPE) at the time of dissection. Each experimental fly was anesthetized on ice for 1 min before being briefly immersed in 70% v/v ethanol to remove food residue and other external debris. Brains were then dissected in phosphate-buffered saline (PBS) solution, before immediately being fixed in 4% v/v paraformaldehyde in PBS for 15 minutes. Fixed tissue was stored at 4º C in PBS for up to 2 weeks before immunostaining.

For larval experiments, brain, ventral nerve cord, and wing discs were collected from 3rd instar wandering larvae. Larval heads were dissected away in PBS and inverted, then immediately fixed in a 4% v/v solution of paraformaldehyde in PBS for 15 min. After fixation, inverted heads were dissected again to extract the specific tissue of interest (brain and nerve cord, or wing discs), which was stored at 4º C in PBS for up to 2 weeks before immunostaining.

Because second generation MCZR constructs were marked only by CyO or Sb, balancers could not be visually confirmed in larvae. We therefore also retained a piece of tissue from the posterior end of each dissected animal, which was stored at -20º C for up to 2 weeks, before it was used for genotyping. DNA was extracted in a 1% v/v solution of Proteinase K in ‘Squishing’ Buffer (0.01M Tris, 0.001M EDTA, 0.025M NaCl) at 37º C for 30 min, followed by a 2 min incubation at 98º C. The following primer pairs were used to detect the presence of genomic Flp or ФC31: Flp_1107F (GAGATGATCGCGCTGAAGGA), Flp_1245R (CGGCGATTGATGTAGGAGCT), ФC31_211F (TCGAACGCATCCTGAACGAA), and ФC31_535R (CGTGATCTCCTTCGTCTCCG). A pair of primers targeting GAL4 were additionally used as a positive control for successful genotyping: Gal4_F (TCTTTGGCGACGGTTCTCTC) and Gal4_R (CCCCCTCTATACACCAGGCT). The following polymerases were used in PCRs: for larval genotyping, Phusion (New England Biolabs, M0530L); for molecular genetics, GoTaq (New England Biolabs, M300F). PCRs were run in a C1000 Touch thermal cycler (BioRad) according to protocols provided by the polymerase manufacturer.

### Immunohistochemistry

All immunohistochemistry experiments were performed according to established protocols. Briefly, fixed tissue was blocked and permeabilized with a 10% v/v solution of normal goat serum in 0.5% v/v PBS-TritonX (PBST), then incubated in primary antibody solution overnight at 4º C on a nutator. Brains were washed three times in PBST, then incubated overnight in secondary antibody solution at 4º C on a nutator. Brains were then washed three more times in PBST, and stored in PBS for up to a week before imaging.

Primary and secondary antibody solutions each contained 5% v/v normal goat serum, and were diluted in PBST. The following primary antibodies were used in this study: mouse anti-NC82/brp (1:25, DSHB #nc82-s), chicken anti-GFP (1:2000, Abcam #13970, lot #GR261775-3), rabbit anti-DsRed (1:2000, Takara #632496, lot #2210019). These primary antibodies were tagged with the following fluorescence-conjugated secondary antibodies: goat anti-mouse/AlexaFluor647 (1:250, Invitrogen #A21240, lot #2482960), goat anti-chicken/AlexaFluor488 (1:250, Invitrogen #11039, lot #2180688), and goat anti-Rabbit/Cy3 (1:250, Invitrogen #A10520, lot #2381135). Larval wing disc experiments (Extended Data Fig. 2) additionally used an AlexaFluor647-conjugated phalloidin counterstain (1:10, Invitrogen #A22287, lot #3140549). All antibodies except mouse anti-NC82 and the phalloidin counterstain were stored as 100x working dilutions in PBS at 4º C, and were routinely evaluated for effectiveness. Undiluted stock solutions for all antibodies were stored according to the manufacturer’s specifications.

### Confocal imaging and asymmetry analysis

Immunostained tissue was immersed in Vectashield Mounting Medium (Vector Laboratories, #H-1000-10) and mounted on slides. Confocal image stacks were collected on a Leica SP8 with standardized imaging parameters and manually inspected in FIJI.

All flies used for MCZR-SPARC asymmetry analyses are listed in Supplementary Table 1 and Supplementary Table 2. For pan-neuronal labeling experiments (Fig. 3A,B and Extended Data Fig. 4B-D), asymmetry was defined as any qualitative (i.e., ON/OFF) left-right difference in effector gene expression between corresponding regions of brain cortex (i.e., cell bodies). More subtle differences in expression levels were not counted as asymmetric. Brains were then classified according to the spatial extent of asymmetric labeling. “Large” asymmetric patches were those that covered at least a quarter of the central brain, while “small” asymmetric patches covered less than a quarter of the central brain. For cell type-specific MCZR-SPARC experiments (Fig. 3C-F and Extended Data Fig. 4E), “complete” asymmetry was defined as all cells of a given type showing effector expression in one hemisphere, and no cells of that type showing expression in the other hemisphere. Brains with expression patterns that were neither symmetric nor completely asymmetric were defined as showing “partial” asymmetry – typically, “partial” asymmetry meant that one or two cells were ON in an otherwise OFF hemisphere, or that one or two cells were OFF in an otherwise ON hemisphere.

### Behavioral testing and analysis

All behavioral experiments were conducted with PiVR^63^. Briefly, the setup comprises a camera directed at a platform which is illuminated from below with infrared (825nm) LEDs. Optogenetic activation was achieved with a second set of red (625nm) LEDs. A “FlyBowl”-inspired behavioral chamber^62^ was placed on top of the illuminated platform. The chamber was 3D printed (Phrozen, Sonic Mini 8K) using a transparent resin (Phrozen, Aqua Clear Plus), then washed and cured according to manufacturer instructions. Chamber design files will be made available on Zenodo. Each chamber was covered with a 4” x 4” sheet of borosilicate glass (McMaster-Carr, #8476K14), which was siliconized with Sigmacote (Millipore Sigma, #SL2-25ML), effectively eliminating “upside-down” walking on the glass surface.

All behavioral testing was conducted in a darkened room at 25º C and 66% relative humidity. For each experiment, 9-13 DPE female flies were immobilized in their food vial on ice and transferred individually into numbered behavior chambers. After a 30 min acclimatization period, behavioral chambers were placed on a light-protected PiVR platform for behavioral tracking at 25 fps. The stimulus protocol consisted of a 2 sec stimulus of 30 μW/mm^2^ red light followed by 10 sec of darkness that served as the inter-stimulus interval. This protocol was repeated continuously for 40 minutes. After the experiment, flies were immobilized by placing the behavior chamber into a freezer. Immobilized flies were stored individually in food vials until dissection, which was performed as described above (“Tissue collection and genotyping”). Dissected brains were immediately mounted on slides in PBS and imaged *ex vivo* using a Leica SP8 confocal microscope, allowing us to confirm the expression pattern of each behaviorally tested fly.

Behavior videos were tracked and analyzed using custom scripts that will be made available on Zenodo. Each video was converted to animal position data using the PiVR tracking algorithm^63^. Positional data was then interpolated and filtered using a Savitzky-Golay filter of length 0.2 sec and a polynomial of order 4. Finally, translational and rotational velocities were calculated as positional differences between timesteps. Flies with an overall mean velocity of less than 2 mm/s were excluded from further analysis. Flies were grouped according to their expression patterns for statistical comparison. “Relative” forward velocities were calculated by subtracting the cross-fly mean “no expression” forward velocity from each individual fly’s cross-trial mean forward velocity (Extended Data Fig. 6A,B). “Sign-corrected” angular velocities for both bilateral and unilateral groups were calculated by multiplying the angular velocity time course of each trial by the sign of the first peak in angular velocity after optogenetic stimulus onset. Cross-trial means for each fly, and cross-fly means for each group, were then calculated as normal. This procedure ensured that the first turn on each trial was positively signed, allowing for the creation and statistical comparison of the unilateral group, which included flies with both “left only” and “right only” expression patterns. Corrected angular accelerations were then calculated as the first derivative of corrected angular velocities.

### 2-photon imaging and analysis

2-photon imaging data was collected on a Bruker Ultima IV with a MaiTai BB femtosecond laser (SpectraPhysics) at 920nm and a Leica 20x HCX APO 1.0 NA water immersion objective. Imaging data was collected at a resolution of 2.6 × 2.6 × 5 μm according to established protocols^15^. Briefly, flies were immobilized on ice and then glued to a custom-cut foil shim. Each fly had the cuticle on the posterior surface of the head dissected away, and was immediately moved to the microscope for imaging. Throughout each experiment, carbogen-bubbled (95% O_2_, 5% CO_2_) imaging saline (103 mM NaCl, 3mM KCl, 5 mM TES, 1 mM NaH2PO4, 4 mM MgCl2, 1.5 mM CaCl2, 10 mM trehalose, 10 mM glucose, 7 mM sucrose, 26 mM NaHCO3) was warmed to 21º C with an in-line heater (Warner Instruments, #SC-20) and continuously perfused across the brain. During imaging, the fly was positioned atop a 9 mm air-suspended polyurethane foam ball (LAST-A-FOAM, #FR4615) so that walking behavior could be tracked as movements of the ball. Ball recordings taken with a Flea FL3-U3-13E4M-c camera were streamed into Fictrac^72^ software which was used to reconstruct a fictive movement trajectory.

All imaging analysis code will be made available on Zenodo. Imaging data was first pre-processed according to established protocols^15^. Briefly, ANTs^73^ was used to motion-correct each brain volume, then bleaching correction was performed by subtracting off a gaussian-filtered (σ = 100 sec) version of the raw fluorescence time course for each voxel. Finally, volumetric ROIs were manually drawn based on mean fluorescence images using Napari^74^.

After pre-processing and ROI selection, we then calculated a fluorescence time course for each ROI. For each z-slice within the volumetric ROI, we defined a baseline fluorescence value (F_q_) as the mean fluorescence over behaviorally quiescent periods (when the fly exhibited less than 0.5 mm/s of forward velocity and less than 10 º/s of rotational velocity for at least 3 sec). We then calculated dF/F_q_ for each time point *t* in the entire fluorescence time course F as:

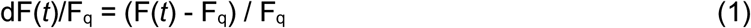

Finally, we pooled across z-slices by summing over voxel count- and sample number-normalized dF/F_q_ time courses for each slice. For plotting (Fig. 5B), we smoothed the z-pooled dF/F_q_ trace with a Savitzky-Golay filter of length 3 sec and a polynomial of order 3. Behavioral traces were similarly smoothed for plotting.

To calculate behavioral filters, we next found the cross-correlation between dF/F_q_ and rotational velocity or forward velocity for each z-slice using the temporal super-resolution method described by Mano et al.^75^. To pool across z-slices, we repeated the same normalize-then-sum process, this time with slice-wise cross-correlations. We then deconvolved the z-pooled cross-correlation *C* with the corresponding behavioral autocorrelation *A*. In the frequency domain, the deconvolved filter *K* was calculated as:

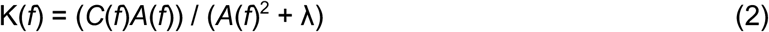

Here λ = 0.001*max(*A*^2^), and acts as a regularization term that prevents the deconvolution division from amplifying noise in frequencies where little signal is present. The filters calculated in this way retained a power peak at exactly the volume imaging rate, which was removed using a notch filter. As a final filtering step, we removed high-frequency ringing with a zero-phase, 4th order Butterworth filter (cutoff = 2.5 Hz). This process produced an angular velocity filter and a forward velocity filter for each ROI in each fly, which we then grouped and averaged according to each fly’s expression pattern. To estimate uncertainty in the cross-fly mean filters shown in Fig. 5, we resampled from the individual filters in each expression group 1000 times (with replacement), allowing us to define a 90% confidence interval.

To quantify the filter difference across brain hemispheres (and thereby compare the relative filter asymmetry for unilateral and bilateral expression groups), we defined a label-agnostic Asymmetry Index (AI). We first found the mean absolute amplitude from time 0 to 0.5 sec for the left and right hemisphere filters for each fly. We next defined the hemisphere with the larger value as K_strong_, and the hemisphere with the smaller value as K_weak_. We then calculated AI as:

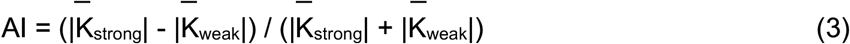

### Connectome analysis

All connectome data included in Extended Data Fig. 5 and Supplementary Table 3 were taken from FAFB dataset^64,76,77^ (ver 783) and analyzed in Codex. 3D reconstructions were generated with Neuroglancer (Google Connectomics Team). The following root IDs were identified as LAL089: 720575940631306148, 720575940624521706, 720575940611790829, 720575940644752494, and 720575940624334606. The following root IDs were identified as LAL091: 720575940637150371, 720575940607480073, 720575940612884433, 720575940631747833, 720575940628656834, and 720575940615482229. The example neuron with annotated synapse positions in Extended Data Fig. 5B is root ID 720575940644752494. When calculating synapse distances from the midline, we estimated its position by taking the mean of brain_mesh_v3. Only synapses with a cleft score greater than 50 were considered in our analysis. When listing top synaptic partners, only cell types that provided/received at least 5% of the *joint* total synapses to/from *both* LAL089 *and* LAL091 were listed^78^.

We note that LAL089 and LAL091 were originally defined in the hemibrain dataset^56^, where the exact number of each type differ from what has been annotated in FAFB (this discrepancy has been previously noted^59^). Our immunohistochemistry experiments indicate that there are 8-9 LAL089/091 neurons per hemisphere, consistent with annotations in the hemibrain^56^ and more recent BANC^65^ datasets. Because only 5-6 cells are labeled per hemisphere in the FAFB dataset, it is likely that some LAL089/091 cells were mislabeled as other cell types. However, it does not appear that non-LAL089/091 cells were mislabeled as LAL089/091. We did not see any evidence that LAL093 (i.e., root ID 720575940625370387) is a part of the *SS49952-GAL4* driver line^59^.

## Large Language Model (LLM) Usage Disclosure

Claude Code with Opus 4.7 and 4.8 (Anthropic) was used in the development of python scripts and marimo notebooks for some elements of the asymmetry rate (Fig. 3) and imaging data (Fig. 5) analysis pipelines. All LLM-generated code was carefully reviewed and tested for accuracy. The authors take full responsibility for the validity of all code used to analyze data and create figures for this work.

## Notes

### Competing Interest Statement

The authors have declared no competing interest.

