## Supplementary Figures for "Unilateral gene expression via maternally controlled zygotic recombination in *Drosophila*"

**List of Supplementary Files**

Supplementary Video 1 – backwards turning with bilateral stimulation of LAL089/091

Supplementary Table 1 – pan-neuronal symmetry scoring data

Supplementary Table 2 – cell type-specific symmetry scoring data

Supplementary Table 3 – top synaptic partners of LAL089/091

Supplementary Table 4 – list of genotypes by figure panel and stock sources

Extended Data Figures 1–7

### Extended Data Figures

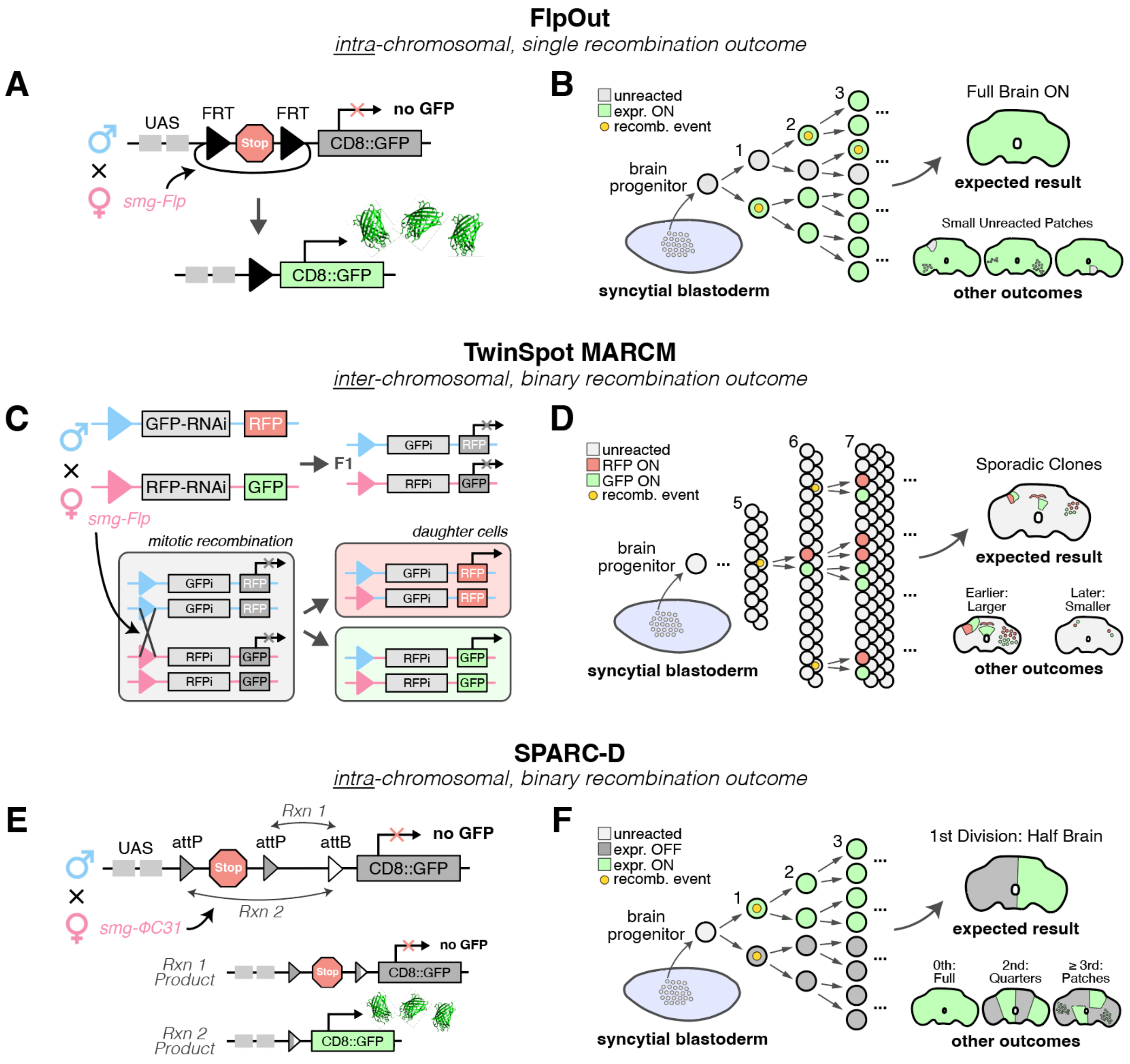


**Extended Data Figure 1.** Details of methods used to assess MCZR minigene activity.
Genetics **(A,C,E)** and expected results **(B,D,F)** are shown for each approach. Numbers in **(B,D,F)** represent the number of times the initial "brain progenitor" has divided, with the brain progenitor defined as the last syncytial nucleus that will give rise to all brain lineages.
**(A)** FlpOut recombination with *smg-Flp* (MCZR-FlpOut). FlpOut relies on an intrachromosomal recombination event that removes a "Stop" cassette comprised of transcriptional and translational terminators[43,44].
**(B)** Because FlpOut recombination only produces one outcome (GFP ON), most or all of the adult brain is expected to be GFP+.
**(C)** TwinSpot MARCM recombination with *smg-Flp* (MCZR-TwinSpot). TwinSpot relies on an interchromosomal recombination event that generates a pair of fluorescent daughter cells from a non-fluorescent mitotic precursor cell[29,45].
**(D)** Because interchromosomal recombination is hypothesized to be less efficient than intrachromosomal, TwinSpot is expected produce sporadic pairs of GFP+ and RFP+ clones in the adult brain, reflecting later recombination events.
**(E)** SPARC-D recombination with *smg-ΦC31* (MCZR-SPARC). SPARC relies on a pair of competing intrachromosomal recombination reactions, with only one reaction removing a "Stop" cassette comprised of transcriptional and translational terminators[46]. See Extended Data Fig. 3 for full example crossing schemes using *smg-ΦC31*.
**(F)** Because SPARC recombination is both two-outcome *and* intrachromosomal, large asymmetric clones are expected to be possible, including clean left-right splits in the adult brain.

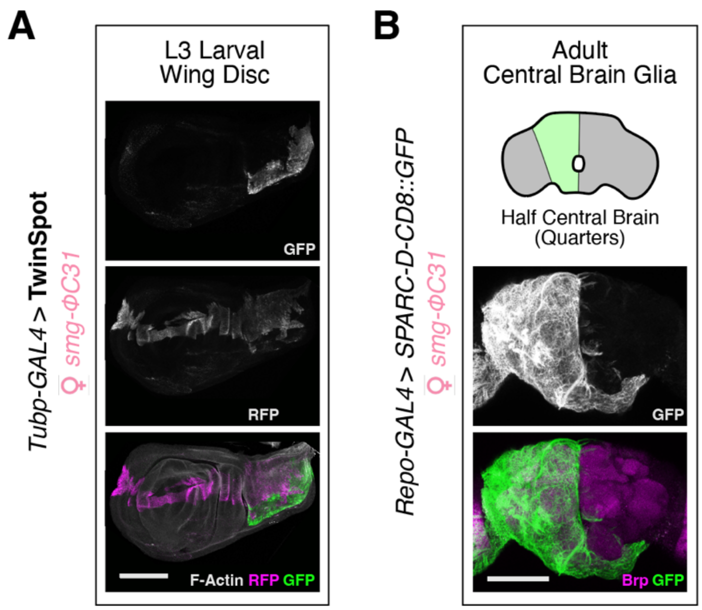


**Extended Data Figure 2.** MCZR functions as expected in non-neuronal cell types.
**(A)** MCZR-TwinSpot in the L3 larval wing disc, plotted as in Fig. 2C,D. F-Actin counterstain shows full disc anatomy. Scale bar: 100 µm.
**(B)** MCZR-SPARC in adult central brain glia, plotted as in Fig. 3A. Brp counterstain shows extent of neuropil. Scale bar: 100 µm.

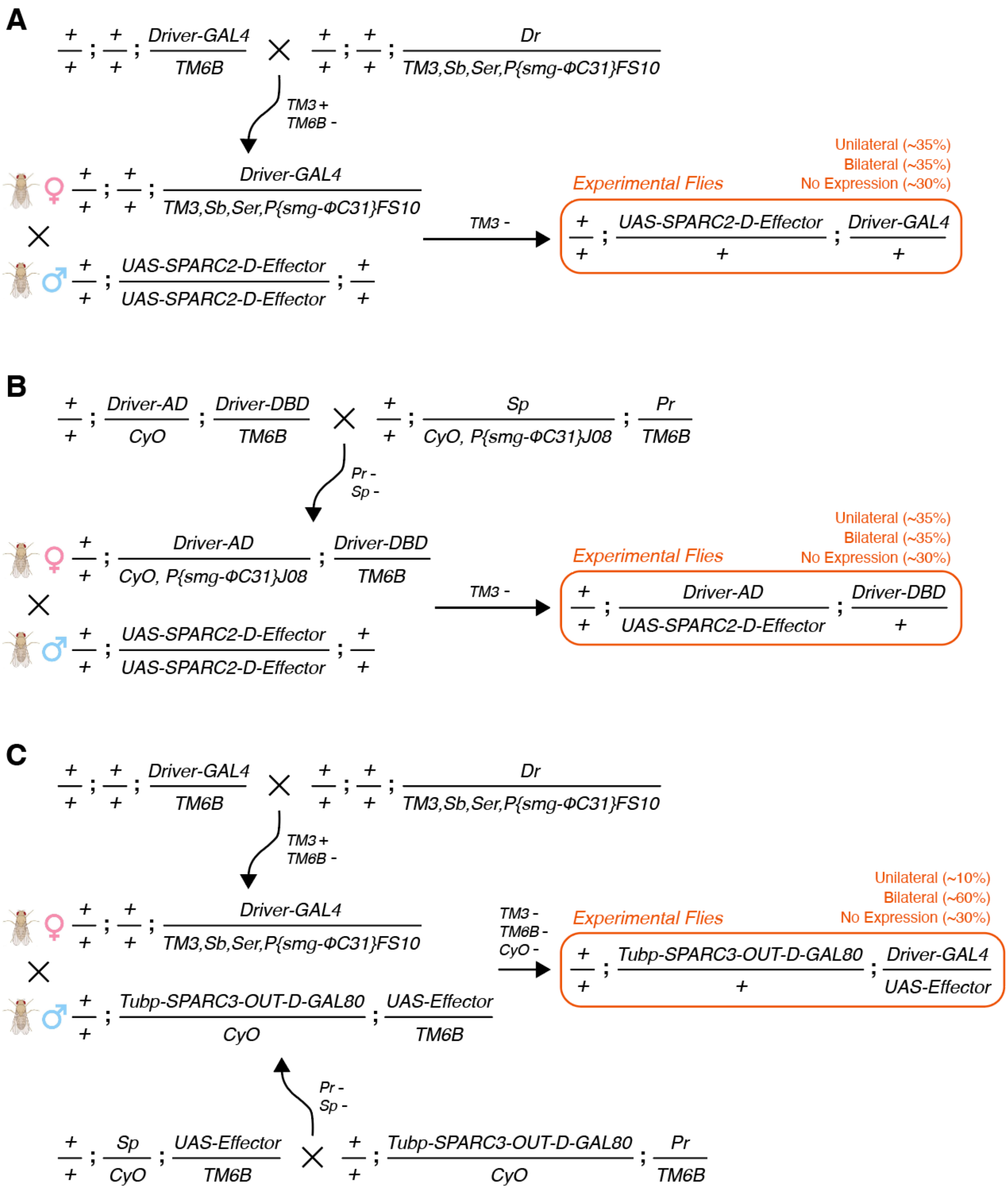


**Extended Data Figure 3.** MCZR-SPARC crossing schemes for unilateral effector expression in a driver line of interest.
**(A)** Crossing scheme for a traditional GAL4 driver line and a SPARC2 effector. The GAL4 line is balanced with *TM3,Sb,Ser,P{smg-ΦC31}FS10*, and virgins of this balanced line are crossed to SPARC males. The recombinase balancer is selected against in the final fly. This cross – and those in (B) and (C) – produces flies with unilateral, bilateral, or no expression in the cells normally targeted by the chosen driver line. Figs. 3A and 6B are examples of this crossing scheme.
**(B)** Crossing scheme for a split-GAL4 driver line and a SPARC2 effector. The split-GAL4 line is balanced with *Sp/CyO,P{smg-ΦC31}J08; Pr/TM6B*, and virgins of this balanced line are crossed to SPARC males. The recombinase balancer and all other balancers are selected against in the final fly. All crosses in Figs. 4 and 5 use this scheme.
**(C)** Crossing scheme for a traditional GAL4 driver line and a traditional UAS effector. The GAL4 line is balanced with *TM3,Sb,Ser,P{smg-ΦC31}FS10*, while the UAS effector is combined with *Tubp-SPARC3-OUT-D-GAL80*. Virgins of the former cross are then mated with males of the latter. The recombinase balancer and all other balancers are selected against in the final fly. Due to the larger inter-recognition site distance in SPARC3-OUT-D, recombination occurs at a lower rate compared to SPARC2-D. Fig. 6A is an example of this crossing scheme.

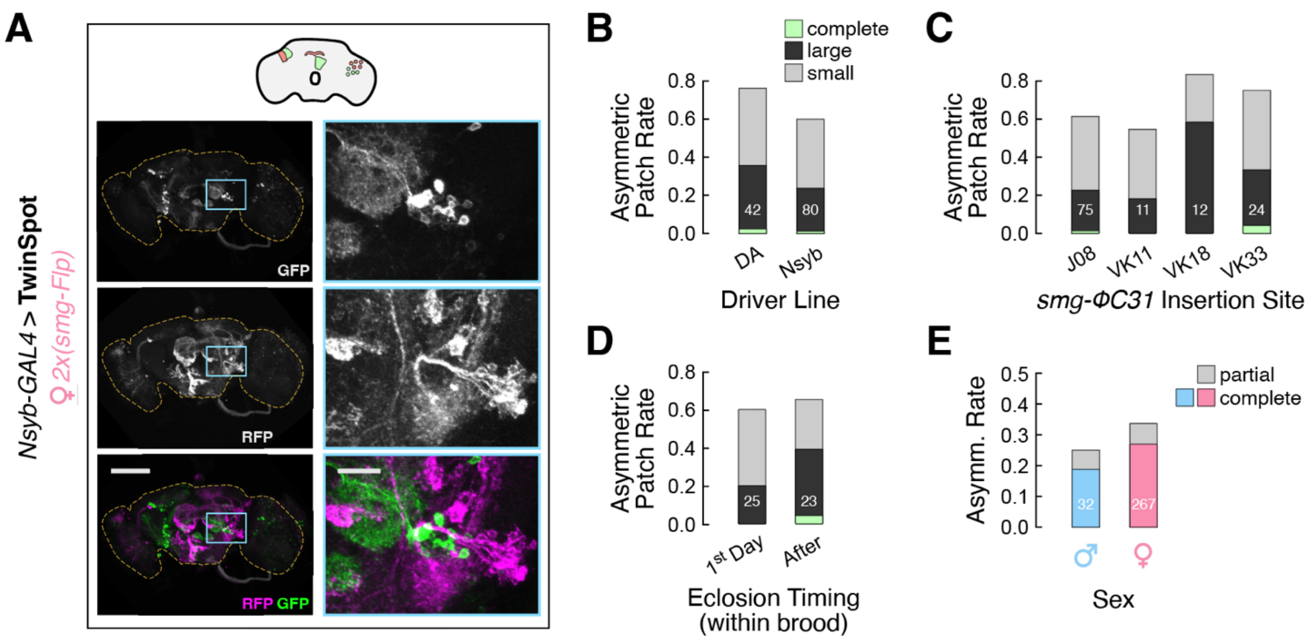


**Extended Data Figure 4.** Many factors do not significantly modulate MCZR function.
**(A)** MCZR-TwinSpot driven by a double genetic dose of maternal *smg-Flp* does not qualitatively alter GFP+ and RFP+ clones, suggesting that mRNA concentration in the mature oocyte is not rate-limiting for *smg-Flp* recombination timing. Plotted as in Fig. 2C,D. Scale bars: 100 µm (left), 20 µm (right).
**(B-D)** Asymmetric GFP+ patch rate by patch size and either **(B)** pan-neuronal driver line, **(C)** *smg-ΦC31* insertion site, or **(D)** offspring eclosion rank within a brood, all for the same data shown in Fig. 3A,B. None of these variables significantly modulate the relative rates of large and small patches (driver: $p$ = 0.54 by Boschloo's Exact, insertion site: $p$ = 0.14 by $\chi^2$ test, eclosion timing: $p$ = 0.16 by Boschloo's Exact). Supplementary Table 1 lists each fly used in these analyses.
**(E)** Sex breakdown of complete and partial asymmetry rates across all cluster 1 cell types (defined in Fig. 3E). Cell types in clusters 2 and 3 were excluded due to their significantly lower complete asymetry rates. Twelve flies from cluster 1 that were missing sex data were also excluded. $p$ = 0.84 by Boschloo's Exact. Supplementary Table 2 lists each fly used in this analysis.

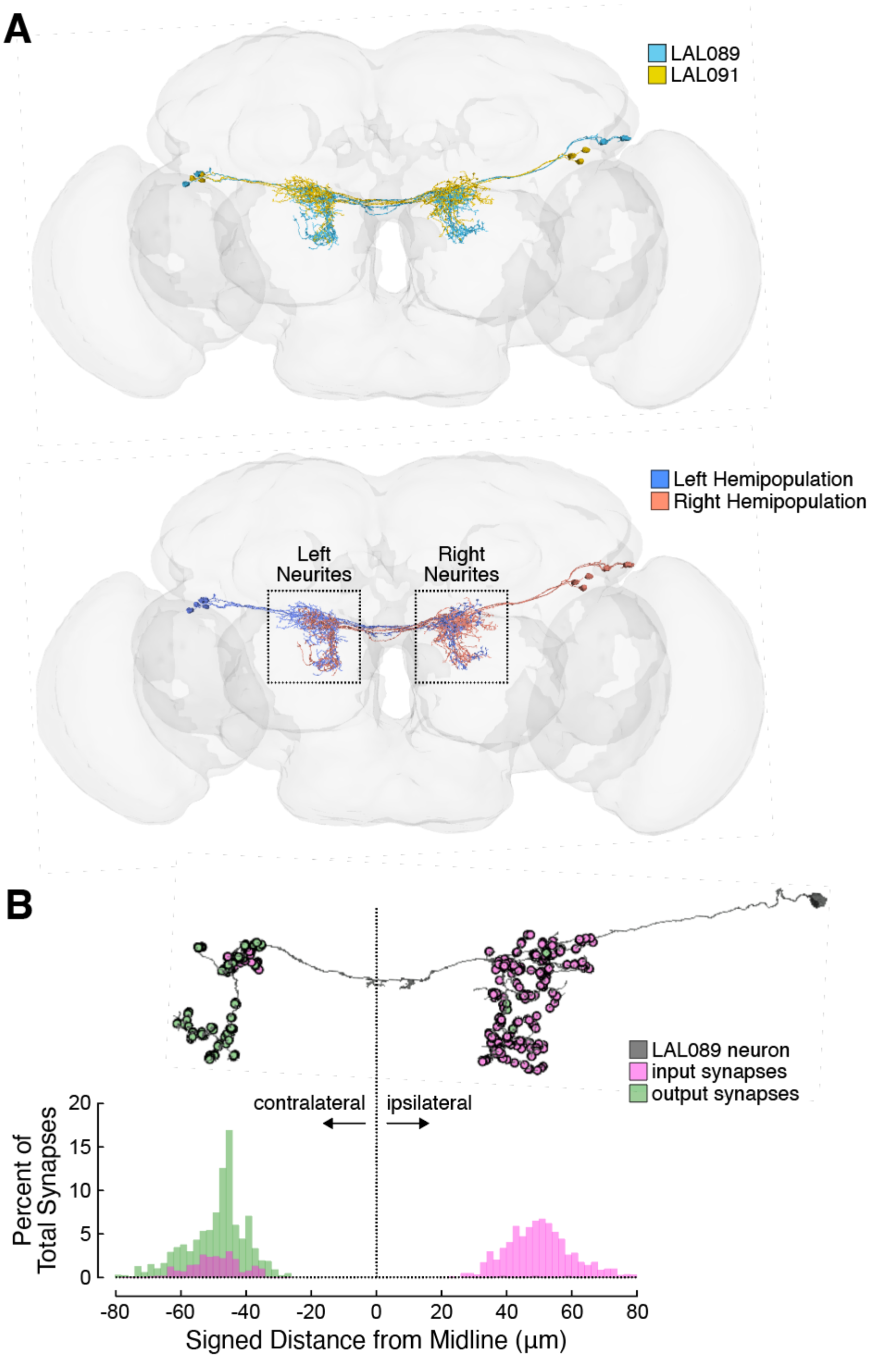


**Extended Data Figure 5.** Connectomic analysis of LAL089 and LAL091.
**(A)** Connectomic reconstructions of LAL089 and LAL091 neurons overlaid on a 3D mesh of the fly brain, using data from the Flywire project[64]. Top: LAL089 and LAL091 are rendered in different colors to highlight structural differences. Bottom: the left and right hemi-populations are rendered in different colors. Neurites in the left and right hemispheres are labeled accordingly – note that "left neurites" and "right neurites" do not mean the same thing as "left hemi-population" and "right hemi-population".
**(B)** Anatomical polarization in LAL089/091. Top: a connectomic reconstruction of a single LAL089 neuron, with pre- and post-synaptic sites colored in green and magenta, respectively. The image is scaled and positioned to match the x-axis below. Bottom: Histogram of synapse positions for all LAL089/091 neurons, relative to the anatomical midline. "Ipsilateral" (positive values) and "contralateral" (negative values) are relative to the hemisphere of each neuron's soma. Percentages shown are for each 2 µm bin, and are calculated independently for inputs ($N$ = 1640) and outputs ($N$ = 2185).

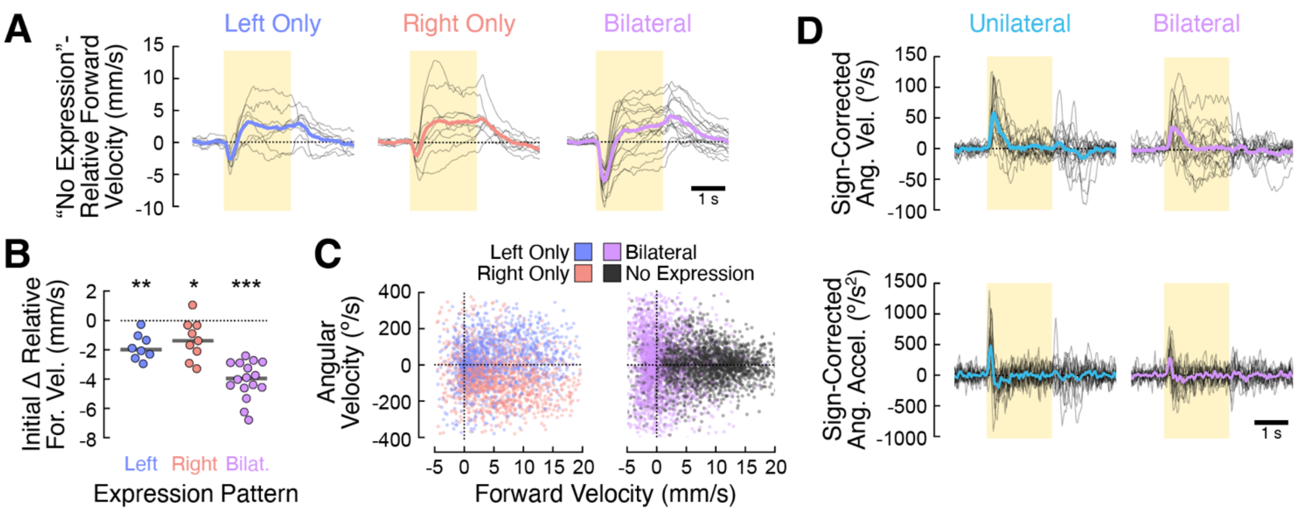


**Extended Data Figure 6.** Additional behavioral analyses.

**(A)** Corrected forward velocity for flies with left- (blue), right- (red), and bilaterally (purple) stimulated LAL089/091 neurons. Each line represents the mean forward velocity of one fly, with both its mean pre-stimulus velocity *and* the cross-fly mean “No Expression” velocity time course subtracted out (Fig. 4D, bottom right). Because “No Expression” flies respond to the red light of the optogenetic stimulus with an increase in forward walking, these traces are closer to the “true” LAL089/091 activation-evoked response for each group.

**(B)** Swarm plots of transient changes to corrected forward velocity as a function of expression pattern, plotted as in Fig. 4E. Compared to the “No expression” group, flies in all other expression pattern groups significantly reduce their forward velocity at the onset of the optogenetic stimulus (left: $p = 0.001$, right: $p = 0.021$, bilateral: $p < 0.001$ by one-sample *t*-test with Bonferroni correction). However, bilateral flies reduce their forward velocity much more than the left and right groups ($p < 0.001$ by one-way ANOVA with Tukey post-hoc test), to the point of even walking backwards, as shown in (C).

**(C)** Scatter plots of angular velocity vs. forward velocity for LAL089/091 > *SPARC-D-CsChrimson* flies, highlighting the backwards turns elicited by bilateral optogenetic stimulation (purple). In contrast, unilaterally stimulated flies (blue and red) generally turn while moving forward. Data are from 0.05 to 0.3 s after the onset of optogenetic activation, as in Fig. 4F. Each point represents one time step for one fly, and data are split between two axes for clarity.

**(D)** Sign-corrected angular velocity (top) and acceleration (bottom) for all flies with unilateral (teal) and bilateral (purple) CsChrimson expression in LAL089/091 neurons, calculated and plotted as in Fig. 4G,H.

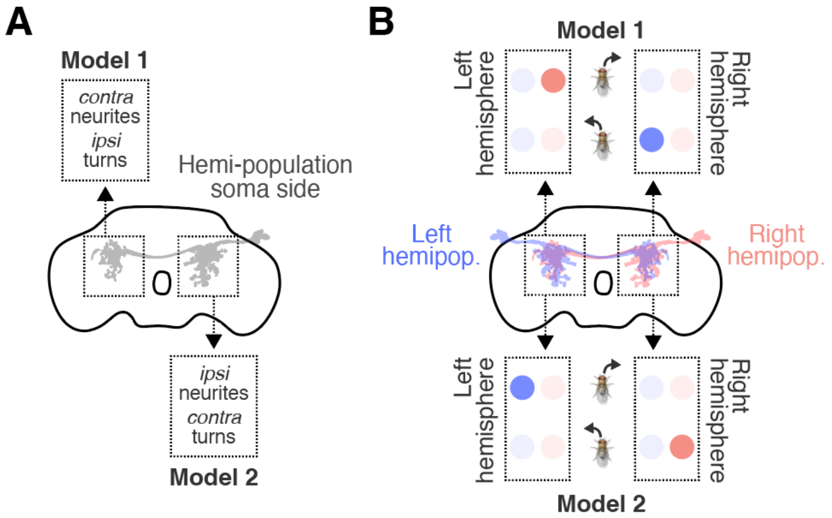


**Extended Data Figure 7.** Models of LAL089/091 locomotor signal encoding.
**(A)** For a given LAL089/091 hemi-population (the set of LAL089/091 neurons with cells bodies in one hemisphere), two encoding models are consistent with our bilateral imaging data (Fig. 5). In one model, ipsilateral turns are encoded in the LAL089/091 neurites contralateral to the cell bodies ("model 1"). In the second model, contralateral turns are encoded in the LAL089/091 neurites ipsilateral to the cell bodies. Note that all uses of "ipsi" and "contra" refer to the hemi-population's side (i.e., cell body location).
**(B)** Models 1 and 2 predict different results for unilateral imaging experiments. The activity of the left and right hemisphere neurites of each LAL089/091 hemi-population is represented as a pair of colored dots, with deeper saturation indicating higher activity levels. Top: model 1 predicts that the left hemi-population will encode left turns in the right hemisphere neurites, and that the right hemi-population will encode right turns in the left hemisphere neurites. This is the model that was confirmed by our data (see Fig. 5C,E). Bottom: model 2 predicts that the left hemi-population will encode right turns in the left hemisphere neurites, and that the right hemi-population will encode left turns in the right hemisphere neurites. This model is not supported by our data.
